# GABAergic neuron dysregulation in a human neurodevelopmental model for major psychiatric disorders

**DOI:** 10.1101/2024.09.23.614564

**Authors:** Ziyuan Guo, Yijing Su, Wei-kai Huang, Xiang Sean Yao, Yan Hong, Alice Gordin, Ha Nam Nguyen, Zhexing Wen, Francisca Rojas Ringeling, Gong Chen, Shiying Li, Lu Lu, Menghang Xia, Wei Zheng, Akira Sawa, Guang Chen, Kimberley M. Christian, Hongjun Song, Guo-li Ming

## Abstract

“GABA dysfunction” is a major hypothesis for the biological basis of schizophrenia with indirect supporting evidence from human post-mortem brain and genetic studies. Patient-derived induced pluripotent stem cells (iPSCs) have emerged as a valuable platform for modeling psychiatric disorders, and previous modeling has revealed glutamatergic synapse deficits. Whether GABAergic synapse properties are affected in patient-derived human neurons and how this impacts neuronal network activity remain poorly understood. Here we optimized a protocol to differentiate iPSCs into highly enriched ganglionic eminence-like neural progenitors and GABAergic neurons. Using a collection of iPSCs derived from patients of psychiatric disorders carrying a *Disrupted-in-Schizophrenia 1* (*DISC1*) mutation and their unaffected family member, together with respective isogenic lines, we identified mutation-dependent deficits in GABAergic synapse formation and function, a phenotype similar to that of mutant glutamatergic neurons. However, mutant glutamatergic and GABAergic neurons contribute differentially to neuronal network excitability and synchrony deficits. Finally, we showed that GABAergic synaptic transmission is also defective in neurons derived from several idiopathic schizophrenia patient iPSCs. Transcriptome analysis further showed some shared gene expression dysregulation, which is more prominent in *DISC1* mutant neurons. Together, our study supports a functional GABAergic synaptic deficit in major psychiatric disorders.

## Introduction

Schizophrenia (SCZ) is a severe psychiatric disorder affecting approximately 1% of the global population, imposing a significant burden on both patients and society^1,2^. While antipsychotic medications exist to relieve symptoms, they fail to address negative and cognitive symptoms and often cause serious side effects^3–5^, indicating that key therapeutic targets may be overlooked. Given the debilitating nature of this disorder and its substantial health and economic impacts, understanding the pathogenic mechanisms underlying SCZ is crucial for developing novel treatments and improving patient outcomes.

There are three major hypotheses of SCZ: genetic risk, aberrant neural development, and synaptic dysregulation^6–10^. Given its high heritability, genetic variants are critical in SCZ’s etiology. Three large genome-wide association studies (GWAS) identified nearly three hundred risk loci linked to SCZ^11–13^, including rare coding variants^13^. However, the pathways through which these loci contribute to disease remain elusive. Much of our current knowledge comes from postmortem immunohistology and transcriptome studies, often confounded by secondary effects from prolonged psychotropic drug use or chronic illness. Therefore, these studies are not ideal for investigating underlying mechanisms. Patient-derived induced pluripotent stem cells (iPSCs) present a valuable model to overcome these limitations, especially during early developmental periods when psychiatric disorders typically emerge. By generating patient-specific brain cells, iPSCs allow researchers to explore the roles of genetic risk factors and uncover molecular and cellular deficits in major psychiatric disorders within the same genetic background as patient brains^14–17^, and previous modeling has revealed glutamatergic synapse deficits^18–24^. Consequently, iPSC models have the potential to bridge the three primary SCZ hypotheses, offering novel insights into disease mechanisms and revealing new therapeutic targets.

Cortical GABAergic interneurons, particularly those derived from the medial ganglionic eminence (MGE), are consistently implicated in SCZ, with supporting evidence from human post-mortem and genetic studies^25–30^. Dysregulation of GABAergic transmission, such as reductions in glutamate decarboxylase 67 (GAD67) and alterations in presynaptic GABA transporters and postsynaptic GABA_A_ receptors, is thought to contribute to cognitive deficits in SCZ by disrupting cortical gamma oscillations^31,32^. An iPSC study found reduced GAD67 and synaptic proteins like gephyrin and neuroligin-2 (NLGN2) in SCZ patient-derived GABAergic neurons^33^. However, it remains unclear how SCZ risk genes causally affect GABAergic synaptic structure and function during early development, whether these phenotypes are present in idiopathic cases, and how dysregulated inhibitory transmission alters cortical neural circuitry in the disease.

In this study, we utilized a collection of iPSC lines from a family with major psychiatric disorders linked to a frameshift mutation in *Disrupted in Schizophrenia 1* (DISC1; D2 and D3) and an unaffected family member with wild-type *DISC1* (C3)^34,35^. We generated isogenic iPSC lines to either rescue the *DISC1* mutation in patient lines (D2R and D3R) or introduce the mutation into the unaffected family member (C3M)^22,23^ (Extended Data Fig. 1a-b and Extended Data Table 1). Our study revealed mutation-dependent deficits in GABAergic synapse formation and function, confirmed in homogenous MGE-derived GABAergic neurons, indicating a cell-type-specific defect in *DISC1* mutant GABAergic neurons. We reconstructed cortical circuitry by co-culturing glutamatergic and GABAergic neurons at a physiological ratio and observed that mutant glutamatergic neurons reduced network activity, while mutant GABAergic neurons disrupted network synchrony. Bulk sequencing of MGE-derived GABAergic neurons highlighted dysregulated genes associated with nervous system development and synaptic transmission. Additionally, idiopathic SCZ lines showed similar alterations in GABAergic transmission and gene expression, suggesting common molecular and cellular pathways between rare and idiopathic SCZ cases.

Intriguingly, the effectiveness of differentially expressed genes (DEGs) in idiopathic GABAergic neurons is lower than in *DISC1* mutant lines, reinforcing the idea that mutant lines may provide a more effective model for studying the disease. Finally, when comparing our data with a human postmortem database, we found that the overlapping DEGs are significantly more enriched in SCZ and other mental disorders than in other diseases. These findings suggest that our model could be valuable for shedding light on the underlying mechanisms of SCZ pathogenesis and identifying potential novel therapeutic targets.

## Results

### Developmental defects in GABAergic synapses of *DISC1* mutant human forebrain neurons

We previously established a protocol to differentiate human iPSCs into forebrain-specific neural progenitors expressing PAX6 and OTX2, which subsequently matured into cortical neurons, consisting of 90% CAMKII^+^ glutamatergic neurons and 10% GAD67^+^ GABAergic neurons^22^ (Extended Data Fig. 1c-h). Through this approach, we identified a *DISC1* mutation-dependent decrease in the frequency, but not the amplitude, of glutamatergic spontaneous synaptic currents (SSCs)^22^. Given the presence of 10% GABAergic neurons in these cultures, we further explored the impact of the *DISC1* mutation on GABAergic synapses.

Starting with two pairs of isogenic lines, we performed immunocytochemistry on 6-week-old parallel cultures at the same neuronal density. We observed reduced densities of GAD65^+^ or vGAT^+^ synaptic puncta in mutant neurons (C3M and D3) compared to their isogenic wild-type (WT) counterparts (C3-1 and D3R), suggesting deficits in GABAergic synapse formation (Fig. 1a-d). To further validate this observation, we conducted whole-cell electrophysiological recordings of GABAergic SSCs from neurons differentiated from three pairs of isogenic lines in parallel cultures with similar neuronal density. Both the frequency and amplitude of GABAergic SSCs were reduced in mutant neurons (C3M, D2, D3) compared to their isogenic WT neurons (C3, D2R, and D3R) in both 4- and 6-week-old cultures (Fig. 1e-g and Extended Data Fig. 2). This indicates that the *DISC1* mutation compromises GABAergic synapse function, a finding further supported by reduced responses to GABA puffing, which suggests a post-synaptic contribution to the defects observed in GABAergic synapses onto primarily glutamatergic neurons (Fig. 1h-i).

**Fig 1.**
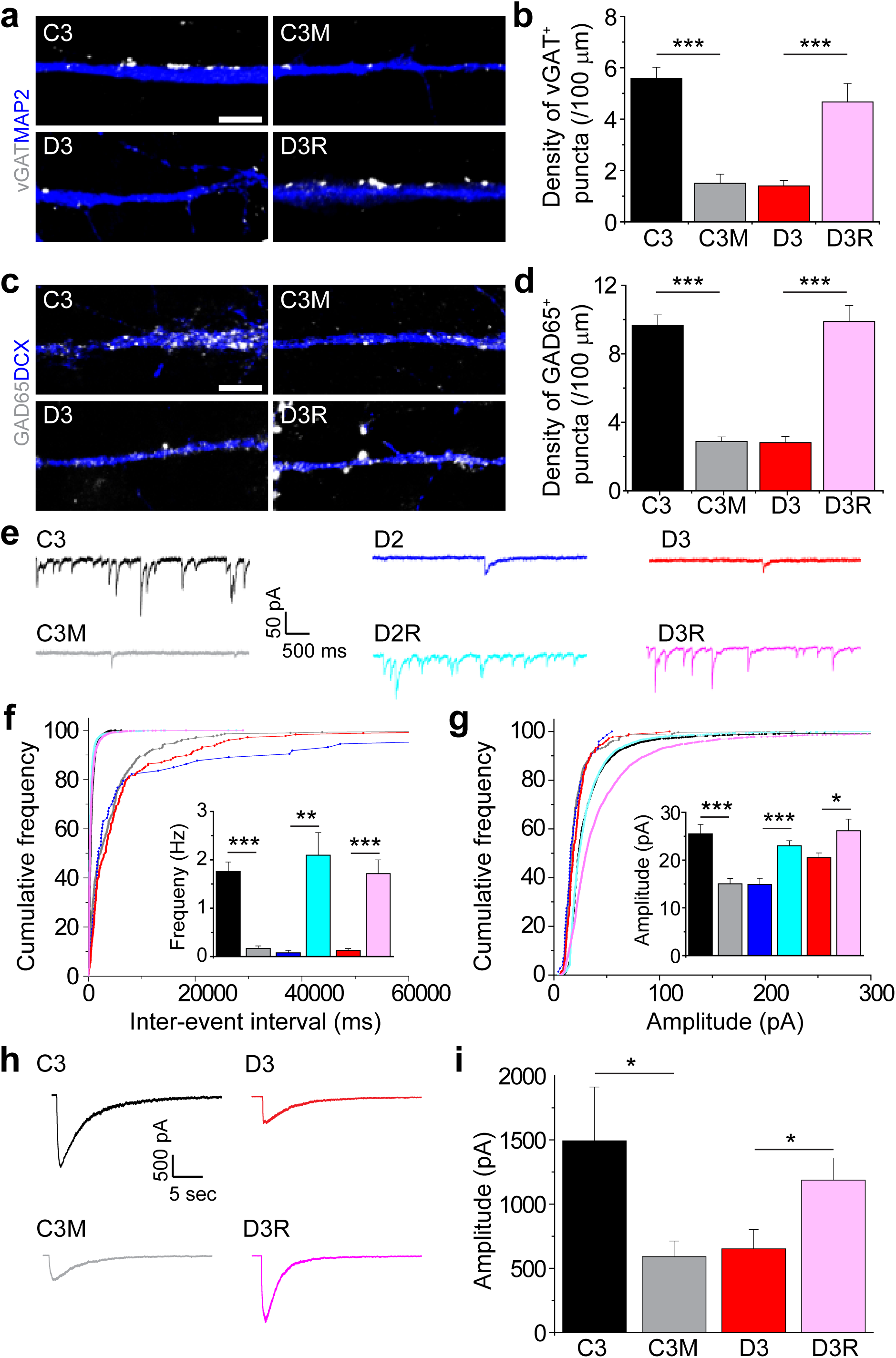
Developmental deficits of GABAergic synapses in human forebrain neurons carrying the *DISC1* mutation. **a-d**, Decreased density of GAD65^+^ and vGAT^+^ puncta in human forebrain neurons carrying the *DISC1* mutation in comparison to isogenic control neurons. Shown are sample images of vGAT (white) immunostaining together with MAP2 (blue) at 4 weeks (**a**) and GAD65 (white) together with DCX (blue) at 6 weeks (**c**) of human neurons after neuronal differentiation (scale bars, 5 µm). Also shown are quantification of vGAT^+^ (**b**) and GAD65^+^ (**d**) puncta density (n = 3 to 11 cultures; data presented as mean ± s.e.m.; ***p < 0.001; Student’s t-test). **e-g**, Defects in GABAergic synaptic transmission in *DISC1* mutant neurons compared to isogenic control neurons. Shown in (**e**) are sample whole-cell voltage-clamp recording traces of spontaneous inhibitory post synaptic currents (IPSCs) of 6-weeks old forebrain neurons in the presence of NBQX (10 µM) and D-AP5 (50 µM). Also shown are cumulative distribution plots of spontaneous IPSCs intervals (**f**) and amplitudes (**g**) (n = 5 to 7 neurons for each condition; Kolmogorov–Smirnov test). Mean frequencies and amplitudes are also shown in the inset (**f, g**). Values represent mean ± s.e.m. (*p < 0.05, **p < 0.01, ***p < 0.001; Student’s t-test). **h-i**, Downregulation of GABA induced currents by *DISC1* mutant neurons. Shown in (**h**) are sample current traces of control and *DISC1* mutant forebrain neurons in response to focal application of GABA (100 µM). Shown in (**i**) is the quantification of GABA induced currents. Values represent mean ± s.e.m. (n = 4 to 8 neurons for each condition; *p < 0.05; Student’s t-test).

### Dysfunction of ganglionic eminence-derived GABAergic neurons due to *DISC1* mutation

During normal human cortical development, most GABAergic neurons originate from the ganglionic eminences^36,37^. To specifically investigate the impact of the *DISC1* mutation on these neurons, we optimized a protocol^38^ to differentiate iPSCs into ganglionic eminence-like neural progenitors, expressing over 90% FOXG1 and NKX2.1 (Extended Data Fig. 3), and then into GAD67^+^ GABAergic neurons (over 90%; Fig. 2a-c). We observed no differences in differentiation between mutant and isogenic WT iPSC lines (Fig. 2b-c and Extended Data Fig. 3b-e).

**Fig 2.**
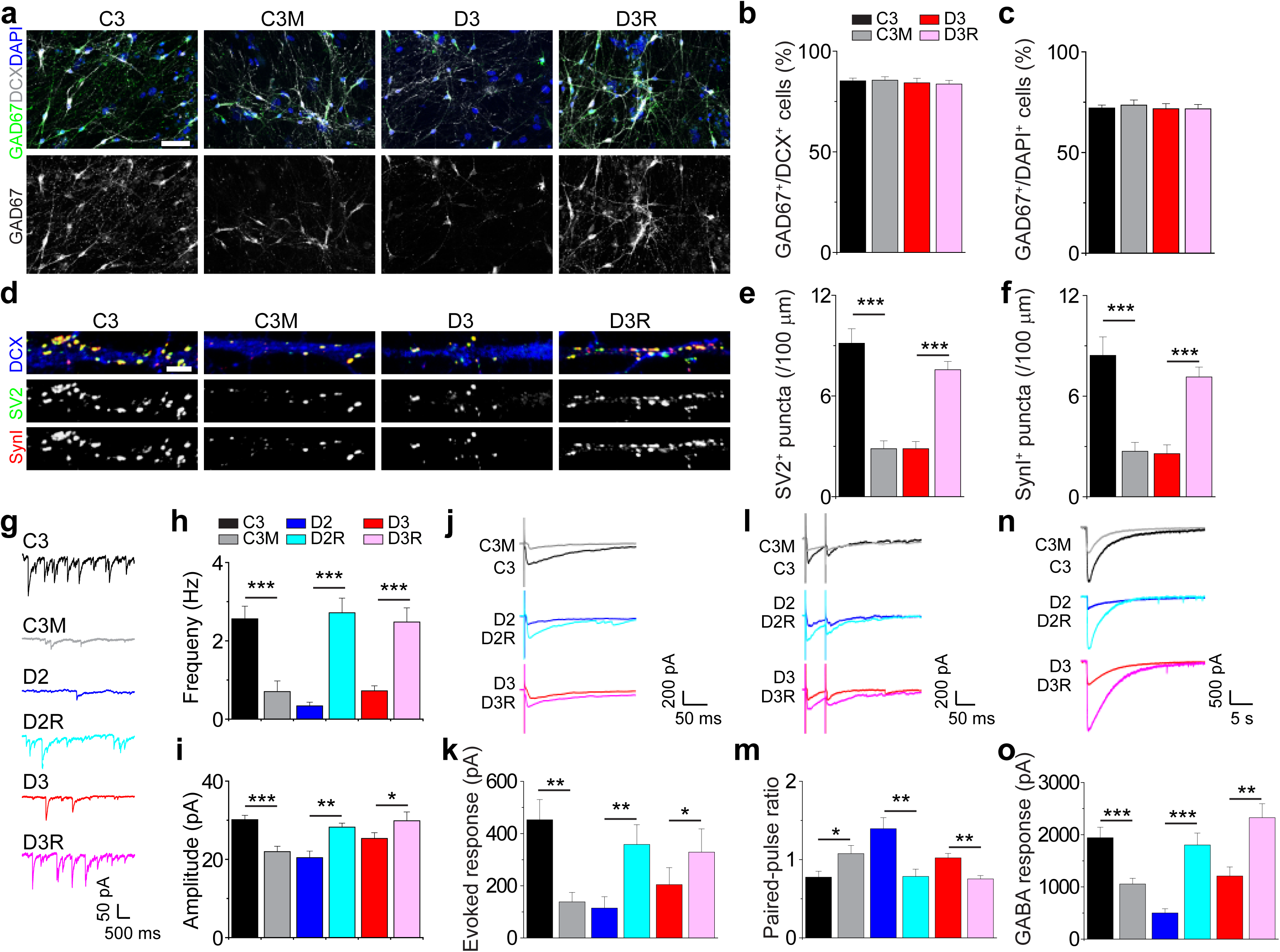
*DISC1* mutant GABAergic neurons display impaired synaptic transmission. **a-c**, Differentiation of iPSCs into GABAergic neurons. Shown in (**a**) are sample confocal images of immunostaining of human GABAergic neurons at 6 weeks after neuronal differentiation for GAD67 and DCX (scale bar, 40 µm). Shown in (**b, c**) are quantifications of GAD67^+^ neurons among different iPSC lines. Values represent mean ± s.e.m. (n = 3-15 cultures). **d-f**, Decreased density of SV2^+^ and SYN1^+^ puncta in human GABAergic neurons carrying the *DISC1* mutation compared to isogenic control neurons. Shown in (**d**) are sample confocal images of SV2 (green) and SYN1 (red) immunostaining (scale bar, 5 µm). Also shown are quantifications of densities of SV2^+^ (**e**) or SYN1^+^ (**f**) puncta of 6-weeks old GABAergic neurons. Values represent mean ± s.e.m. (n = 7 cultures; ***p < 0.001; Student’s t-test). **g-i**, Defects in spontaneous synaptic transmission in *DISC* mutant GABAergic neurons in comparison to isogenic control neurons. Shown in (**g**) are sample traces of spontaneous IPSCs of 6-weeks old GABAergic neurons. Also shown are quantification of mean frequencies (**h**) and amplitudes (**i**) of spontaneous IPSCs. Values represent mean ± s.e.m. (n = 6 to 29 neurons for each condition; *p < 0.05, **p < 0.01, ***p < 0.001; Student’s t-test). **j-k**, Defects in evoked synaptic transmission in *DISC* mutant GABAergic neurons in comparison to isogenic control neurons. Shown in (**j**) are sample traces of evoked IPSCs of 6-weeks old GABAergic neurons. Shown in (**k**) is the quantification of evoked IPSCs. Values represent mean ± s.e.m. (n = 5 to 15 neurons for each condition; *p < 0.05, **p < 0.01; Student’s t-test). **l-m**, Altered presynaptic properties of *DISC* mutant GABAergic neurons in comparison to isogenic control neurons. Shown are sample recording traces (**l**) and quantification of pair-pulse ratio (**m**). Values represent mean ± s.e.m. (n = 4 to 9 neurons for each condition; *p < 0.05, **p < 0.01; Student’s t-test). **n-o**, Downregulation of GABA induced currents by *DISC1* mutant neurons. Shown in (**n**) are sample current traces of control and *DISC1* mutant GABAergic neurons in response to focal application of GABA (100 µM). Shown in (**o**) is the quantification of GABA induced currents. Values represent mean ± s.e.m. (n = 8 to 16 neurons for each condition; **p < 0.01, ***p < 0.001; Student’s t-test).

Next, we examined synapse formation in these GABAergic neurons. Quantitative analysis revealed reduced densities of SV2^+^ and Synapsin I^+^ puncta in mutant neurons (C3M and D2) compared to their isogenic WT counterparts (C3 and D3R) in 6-week-old parallel cultures of the same neuronal density (Fig. 2d-f). Given that over 90% of the neurons in these cultures were GABAergic (Fig. 2b), this finding suggests cell-type specific deficits in GABAergic synapse formation. Consistently, the density of vGAT^+^ synaptic puncta was also reduced in mutant neurons (Extended Data Fig. 4a-b).

Electrophysiological recordings demonstrated decreased frequency and amplitude of GABAergic SSCs in mutant neurons (C3M, D2 and D3) compared to isogenic WT neurons (C3, D2R and D3R) in both 6- and 8-week cultures (Fig. 2g-i and Extended Data Fig. 4c-e and Extended Data Fig. 5a-c). We also observed reduced evoked GABAergic synaptic transmission in mutant neurons (Fig. 2j-k and Extended Data Fig. 5d). To explore pre- and post-synaptic contributions to these deficits, we assessed pair-pulse facilitation (PPF), a presynaptic property^39^, and found an increased PPF ratio in mutant neurons in both 6- and 8- week cultures, indicating dysregulation of presynaptic plasticity (Fig. 2l-m and Extended Data Fig. 5e). Additionally, reduced responses to GABA puffing were observed in mutant neurons, suggesting post-synaptic dysfunction (Fig. 2n-o and Extended Data Fig. 5f), which is similar to what we observed for GABAergic synapses onto mostly glutamatergic neurons (Fig. 1h-i). Interestingly, our previous studies identified presynaptic glutamate release deficits in glutamatergic synapses, but not postsynaptic deficits^22^. Thus, the *DISC1* mutation appears to induce presynaptic deficits in both GABAergic and glutamatergic synapses, along with additional postsynaptic deficits specifically in GABAergic synapses.

### Abnormal neuronal network activity and synchrony resulting from *DISC1* mutant glutamatergic and GABAergic neurons

To model cortical neuronal circuit properties *in vitro*, we differentiated iPSC lines into forebrain glutamatergic and ganglionic eminence-like GABAergic neurons, co-culturing them at an 8:2 ratio on a 64-channel micro-electrode array (MEA)^40^ (Extended Data Fig. 6a). Our analysis focused on 6- week-old neurons, when calcium imaging showed that most neurons had shifted their GABA response from depolarization to hyperpolarization, with increased expression of the chloride transporter KCC2^41^ (Extended Data Fig. 7). To dissect the specific effects of mutant glutamatergic and GABAergic neurons on network properties, we examined four combinations: Glu^WT^/GABA^WT^, Glu^WT^/GABA^MUT^, Glu^MUT^/GABA^WT^, and Glu^MUT^/GABA^MUT^ (C3 as WT, D3 as MUT; Fig. 3a). We observed a range of neuronal activity, from individual firing to synchronized bursts across electrodes (Fig. 3a and Extended Data Fig. 6b). The mean firing frequency of individual spikes was higher in Glu^WT^/GABA^WT^ and Glu^WT^/GABA^MUT^ groups compared to Glu^MUT^/GABA^MUT^ and Glu^MUT^/GABA^WT^ groups (Fig. 3b), indicating that mutant glutamatergic neurons significantly reduce individual neuronal firing within the network.

**Fig 3.**
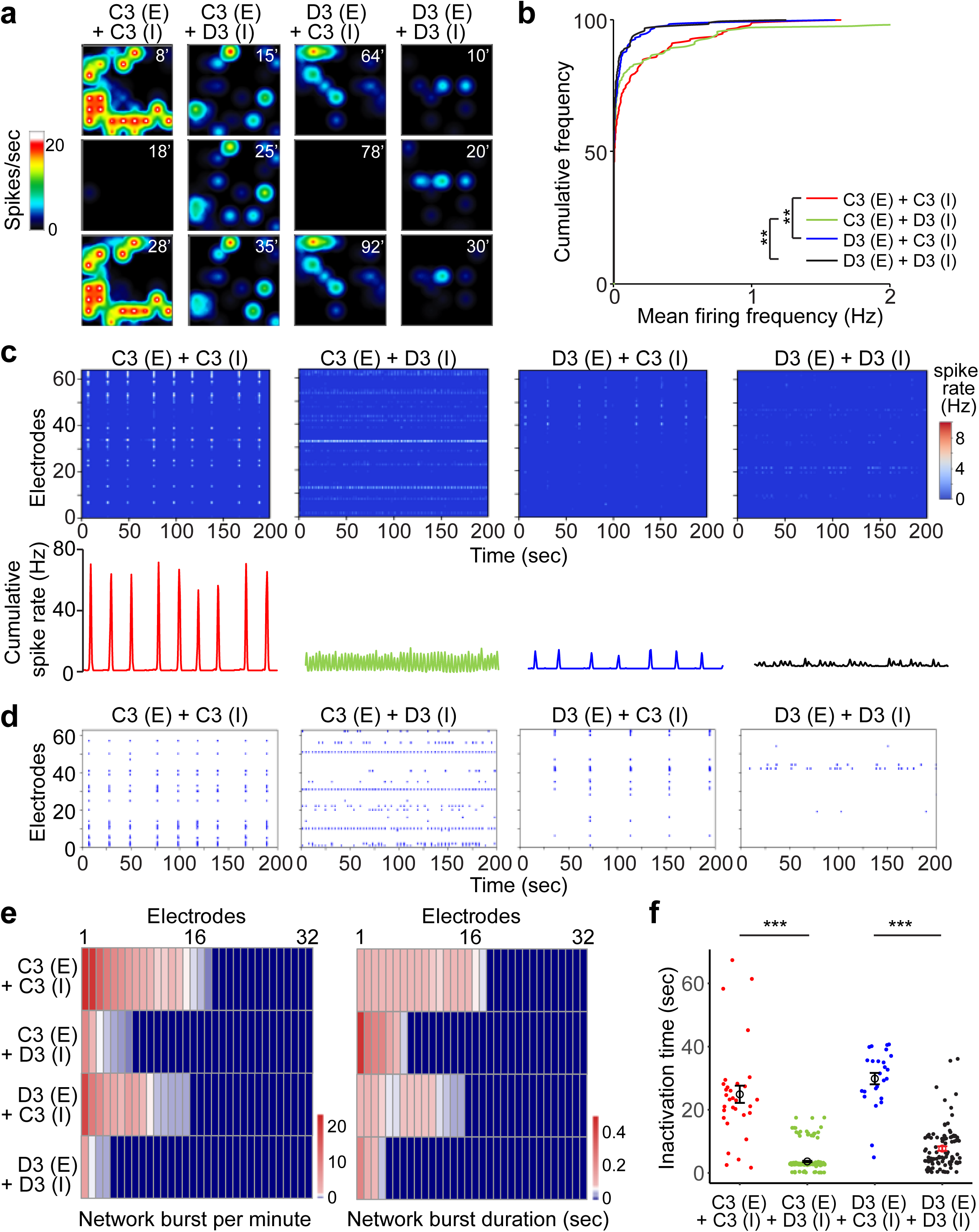
Dysregulation of neuronal network activity and synchronization by *DISC1* mutant glutamatergic and GABAergic neurons. **a**, Real-time heatmaps of spike rate of glutamatergic and GABAergic neuron co-cultures detected at 6 weeks after neuronal differentiation by MEA for four different conditions. **b,** Cumulative distribution plots of mean firing frequency (n = 3 cultures; **P < 0.01; Kolmogorov–Smirnov test). **c,** Sample heatmaps of spike rate and sample cumulative spike rate over 64 electrodes in 200 seconds. **d,** Sample raster maps of bursts over 64 electrodes in 200 seconds. **e,** Quantification of network burst per minute and network burst duration for co-activation of different numbers of electrodes (n = 3 cultures, color codes number of network burst per minute and network burst duration (second) respectively). **f,** Quantification and dot plot of inactivation time between network bursts. Values representing mean ± s.e.m. are also shown (n = 3 cultures; ***p < 0.001; Student’s t-test).

Burst firing analysis revealed higher intra-burst firing frequency but lower burst frequency in Glu^WT^/GABA^WT^ and Glu^MUT^/GABA^WT^ groups compared to Glu^WT^/GABA^MUT^ and Glu^MUT^/GABA^MUT^ groups (Fig. 3c and Extended Data Fig. 6c-f), indicating differences in network properties. Next, we evaluated synchronous firing within the neuronal network, quantifying the degree of synchrony (number of co-active electrodes within a 10 ms window), frequency, and duration. Interestingly, Glu^WT^/GABA^WT^ displayed the highest synchrony degree at higher frequencies, a pattern closely resembling Glu^MUT^/GABA^WT^ (Fig. 3d). Conversely, Glu^WT^/GABA^MUT^ and Glu^MUT^/GABA^MUT^ groups showed reduced synchrony at lower frequencies (Fig. 3d). Similarly, Glu^WT^/GABA^WT^ exhibited longer synchrony durations at higher frequencies, aligning more with Glu^MUT^/GABA^WT^, while Glu^WT^/GABA^MUT^ and Glu^MUT^/GABA^MUT^ showed shorter synchrony durations at lower frequencies (Fig. 3e). Reflecting a more synchronized network, the inactivation time was longer in Glu^WT^/GABA^WT^ and Glu^MUT^/GABA^WT^ compared to Glu^WT^/GABA^MUT^ and Glu^MUT^/GABA^MUT^ groups (Fig. 3f). Collectively, these findings indicate differential contributions of mutant GABAergic and glutamatergic neurons to neuronal network properties.

### Conserved dysregulation of synaptic formation and transmission in glutamatergic and GABAergic neurons from idiopathic SCZ subjects

To further investigate whether dysregulated synaptic transmission is a widespread phenomenon in SCZ neurons, we generated iPSC lines from three idiopathic SCZ patients (SZ49, SZ59, SZ69) and three matched controls (C65, C67, C75) (Extended data Fig. 8 and Extended Data Table 1). After differentiating these iPSCs into forebrain cortical neurons, we observed that all six lines exhibited comparable neural differentiation efficacy (Extended Data Fig. 9). Immunostaining showed approximately 90% of neurons were vGLUT1^+^ glutamatergic and 10% were GAD67^+^ GABAergic across all lines (Extended Data Fig. 10), similar to the *DISC1* cohort^22^. Electrophysiological recordings revealed a significant reduction in the frequency—but not the amplitude—of glutamatergic SSCs in SCZ-derived neurons (Extended Data Fig. 11a-c), consistent with observations in *DISC1* mutant neurons^22^. Additionally, GABAergic transmission was similarly affected in SCZ-derived neurons, with reduced frequency and amplitude of GABAergic SSCs compared to controls (Extended Data Fig. 11d-f). This pattern mirrors the deficits observed in *DISC1* mutant neurons (Fig. 1e-g and Extended Data Fig. 2), indicating that synaptic transmission dysfunction is not unique to the *DISC1* mutation but may be a common feature of SCZ pathophysiology.

To delve deeper, we differentiated the iPSC lines into ganglionic eminence-like neural progenitors and subsequently GABAergic neurons. Immunostaining showed no difference in the efficiency of GABAergic differentiation between SCZ and control lines (Extended Data Fig. 12). However, SCZ-derived GABAergic neurons displayed reduced synaptic puncta density (Fig. 4a-b), marked by lower SV2^+^ staining, indicating impaired GABAergic synapse formation.

**Fig 4.**
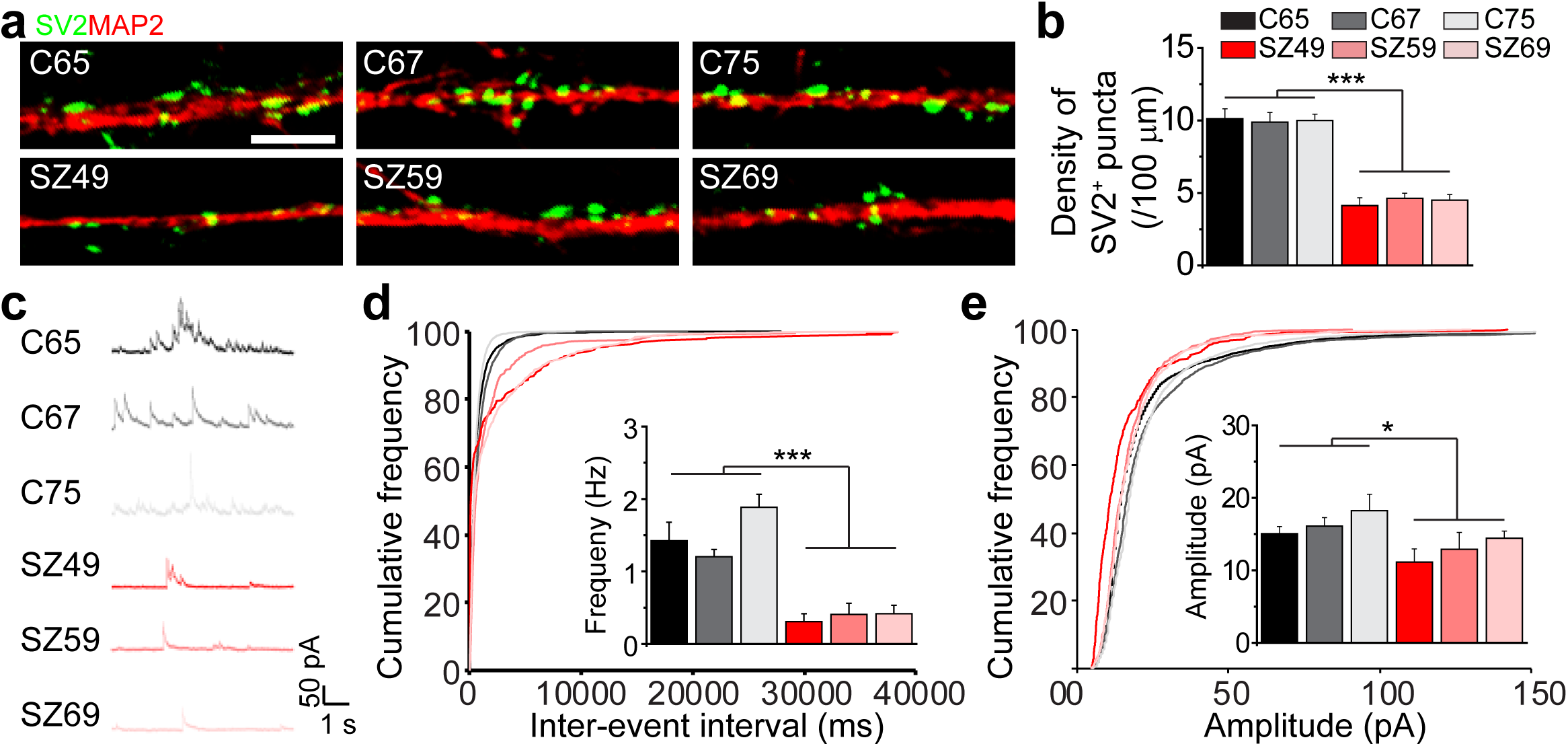
GABAergic neurons derived from idiopathic schizophrenia display impaired synaptic transmission. **a-b**, Reduced density of SV2^+^ puncta in GABAergic neurons derived from idiopathic schizophrenia patient iPSC lines compared to matched control iPSC lines. Sown in (**a**) are sample images of SV2 (green) together with MAP2 (red) immunostaining of human neurons at 6 weeks after GABAergic neuronal differentiation (scale bar, 5 µm). Shown in (**b**) is the quantification of SV2 puncta density. Values represent mean ± s.e.m. (n = 8 cultures; ***p < 0.001; Student’s t-test). **c-e**, Dysregulation of synaptic transmission in ganglionic eminence-derived GABAergic neurons of idiopathic schizophrenia patient iPSC lines in comparison to matched control iPSC lines. Shown in (**c**) are sample recording traces of spontaneous IPSCs of 6-weeks old GABAergic neurons (V_m_ = 20 mV). Also shown are cumulative distribution plots of spontaneous IPSCs intervals (**d**) and amplitudes (**e**). Bar graphs show the mean frequencies and amplitudes of spontaneous IPSCs. Values represent mean ± s.e.m. (n = 6 to 10 neurons for each condition; *p < 0.05, ***p < 0.001; Student’s t-test).

Electrophysiological recordings further confirmed these findings, showing reduced frequency and amplitude of GABAergic SSCs in SCZ neurons (Fig. 4c-e).

Taken together, these results suggest that both glutamatergic and GABAergic synaptic transmission deficits are common features in SCZ neurons, paralleling those observed in *DISC1* mutant neurons. This indicates a shared underlying pathology in both idiopathic and *DISC1*-linked SCZ, providing further evidence that dysregulated synaptic transmission is a core feature of the disease and highlighting the utility of these models for understanding SCZ pathogenesis.

### Transcriptomic dysregulation in *DISC1* mutant and idiopathic schizophrenia-derived GABAergic neurons

Finally, we investigated gene expression changes in GABAergic neurons derived from both the *DISC1* cohort and idiopathic schizophrenia/matched control cohort through transcriptome analysis using RNA sequencing (RNA-seq) on 6-week-old cultures. In the *DISC1* cohort, a common set of differentially expressed genes was identified by comparing two pairs of isogenic lines (C3M vs. C3 and D3 vs. D3R) (Extended Data Fig. 13a-b). Gene ontology (GO) analysis further confirmed that downregulated genes are primarily involved in pathways related to synaptic transmission and neural development (Extended Data Fig. 13c). This finding aligns with the synaptic deficits observed in earlier functional studies. Disease ontology highlighted a significant enrichment of genes linked to schizophrenia (Extended Data Fig. 13d-e). This suggests that DISC1 may serve as a critical regulatory hub in GABAergic neurons, influencing the transcriptional regulation of genes associated with psychiatric disorders.

We extended our analysis to neurons derived from the idiopathic SCZ cohort and their matched controls. Here, GO analysis similarly identified downregulated genes linked to synaptic transmission and neural development (Extended Data Fig. 14a). Disease ontology also pointed to an enrichment of genes related to several psychiatric disorders, including schizophrenia (Extended Data Fig. 14b-c). Importantly, we observed a significant overlap in the sets of dysregulated genes between the DISC1 and idiopathic SCZ cohorts (Fig. 5a-b), with the degree of differential expression being more pronounced in the *DISC1* cohort (Fig. 5c-d). This finding underscores the potential of *DISC1* mutant models to reveal stronger molecular signatures of disease compared to idiopathic cases. Despite the genetic heterogeneity of schizophrenia, these results suggest that there are conserved molecular pathways disrupted in both *DISC1* mutant and idiopathic SCZ neurons, pointing to potential targets for therapeutic intervention.

**Fig 5.**
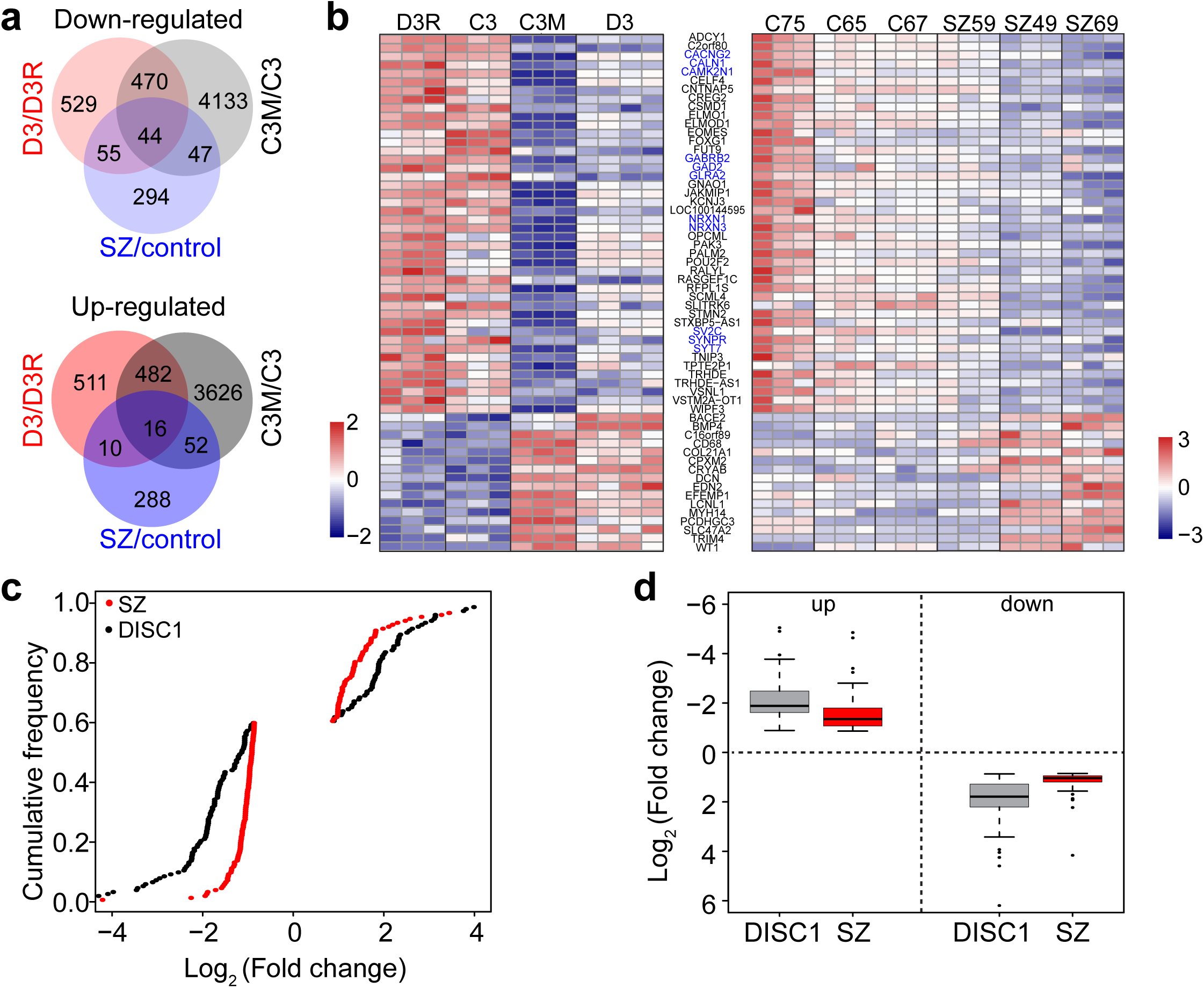
Transcriptomic dysregulation in GABAergic neurons derived from *DISC1* mutant and idiopathic schizophrenia subjects. **a**, Venn diagrams comparing the overlap in differential gene expression between *DISC1* isogenic pairs and idiopathic schizophrenia patients and matched control pairs (n = 3 to 4 samples for each iPSC lines, p value < 0.01 and fold change >1.8; significantly overlapping was tested by fisher test, up regulation p value = 4.5e-3, down regulation p value = 2.3e-17). **b**, Heat map of overlapped differentially expressed genes in the *DISC1* cohort and idiopathic schizophrenia patient/matched control cohort. **c-d**, Decreased efficacy of dysregulated genes in idiopathic schizophrenia/matched control cohort in comparison to the *DISC1* cohort. Shown are cumulative plot (**c**; p value = 0.0009575) and Wiskers plot for the fold change of dysregulated genes in two different cohorts (**d;** up regulation p value = 0.000536, down regulation p value = 1.015e-13).

## Discussion

In this study, we discovered significant dysregulation in GABAergic synaptic formation and transmission in *DISC1* mutant neurons during development. These abnormalities mirror those observed in idiopathic SCZ neurons, indicating that GABAergic synaptic dysregulation is a common cellular phenotype in SCZ. Interestingly, we found that patient-derived glutamatergic and GABAergic neurons have distinct functional roles in cortical circuitry, with *DISC1* mutant glutamatergic neurons reducing network firing and mutant GABAergic neurons disrupting synchronization. RNA-seq analysis revealed shared dysregulation in synaptic transmission and neural development pathways across both *DISC1* and idiopathic SCZ lines, emphasizing the importance of pathway-level understanding in SCZ pathogenesis^42,43^.

Although the ’synaptic dysfunction’ hypothesis has been proposed for many years^8,9,44–46^, it remains unclear how SCZ risk loci influence synaptogenesis and which synapses are affected during early development. Our previous work with iPSC lines from a *DISC1* mutation family^34,35^ revealed deficits in glutamatergic synapse formation and function^22^. This study further shows significant reductions in GABAergic synapse formation and inhibitory transmission using *DISC1* mutant and isogenic control lines, reinforcing the idea that these defects are dependent on the *DISC1* mutation. We generated homogeneous ganglionic eminence-derived GABAergic neurons, observing the same abnormalities without SCZ glutamatergic input, highlighting a cell-type-specific defect in GABAergic transmission during early brain development. Interestingly, both glutamatergic and GABAergic neurons shared presynaptic deficits and postsynaptic response to GABA, but not for postsynaptic response to glutamate. RNA-seq analysis also revealed distinct dysregulated genes in GABAergic neurons compared to the ones in glutamatergic neurons^22^, despite similar GO terms, indicating different molecular mechanisms even within the same risk loci. This underscores the importance of studying homogeneous neuronal populations to better understand SCZ pathogenesis.

Disturbances in working memory are a core feature of schizophrenia, with impaired GABA-mediated network synchronization, especially in the prefrontal cortex (PFC), proposed as a key mechanism^47–49^. Previous iPSC studies often lacked evidence of network alterations. In our study, using a defined co-culture system of forebrain glutamatergic and GABAergic neurons, we demonstrated that *DISC1* mutant glutamatergic neurons exhibit reduced network firing, while mutant GABAergic neurons affect synchronous firing, highlighting cell-type-specific contributions to network dysfunction. These findings provide direct evidence that patient-derived GABAergic neurons cause anomalies in neural synchrony, consistent with abnormal beta- and gamma-band activity in schizophrenia^49,50^. Although our 2D cultures did not replicate these brain waves, suggesting limitations in the model, future use of 3D organoids might capture these dynamics^51–53^. Our results emphasize the potential role of GABAergic dysfunction in the network synchronization and cognitive deficits of schizophrenia, offering insights into novel therapeutic targets at the circuitry level.

The etiology of schizophrenia is highly complex due to its polygenic nature, with numerous risk factors identified through large-scale genome-wide studies^11–13^, including rare coding mutations^13,54^, copy number variants (CNVs)^55^, and common SNPs^11^. Therefore, identifying the common cellular and molecular signatures of polygenic SCZ is crucial for better understanding its etiology. This is especially important given the complex genetics of SCZ, with over two hundred identified risk alleles^11–13^, each likely exerting only a moderate effect, which are challenging to replicate in experimental animals. Although results from post-mortem studies vary, the strongest findings link SCZ-associated variants to synaptic biology pathways^11,12,54,56^. Our study highlights common cellular and molecular signatures of polygenic SCZ by demonstrating GABAergic synaptic dysfunction in both *DISC1* mutant neurons and neurons derived from idiopathic SCZ patients, accompanied by downregulated excitatory transmission^22^, particularly presynaptic vesicle release. These findings suggest that defects in both excitatory and inhibitory synaptic transmission are universal pathological features of SCZ. Additionally, the larger differential gene expression in the *DISC1* cohort may result from the controlled genetic background and DISC1’s role as a hub gene, highlighting the importance of isogenic iPSC studies. Finally, increasing sample size in future studies is crucial for detecting mild SCZ phenotypes.

Together, our study, which examines human GABAergic neurons derived from a variety of patient and isogenic *DISC1* iPSC line pairs, idiopathic schizophrenia iPSC lines, and matched controls, offers a potential mechanistic link among three major hypotheses of schizophrenia and complex psychiatric disorders: genetic risk, aberrant neural development, and GABAergic synapse dysfunction. These findings contribute valuable insights into the general pathogenesis of major psychiatric disorders and establish a foundation for future mechanistic analyses and therapeutic exploration.

## Acknowledgements

We thank member of Ming and Song laboratories for comments and suggestions, D. Johnson, B. Temsamrit, and E. Lanoce for technical support and J. Schnoll for lab coordination. This work was a component of the National Cooperative Reprogrammed Cell Research Groups (NCRCRG) to Study Mental Illness and was supported by the National Institutes of Health (NIH) grant to G-l. M., G.C., and H.S (U19MH106434). Additional supports were from NIH (R01MH105128 and R35NS097370) and Dr. Miriam and Sheldon G. Adelson Medical Research Foundation to G-l.M., and by a postdoctoral fellowship from MSCRF to Z.G., by a Young Investigator Award from NARSAD to Y.S. M.X. and W.Z. were supported by the Intramural Research Program of the National Center for Advancing Translational Sciences.

## Author contributions

Z.G. led and was involved in every aspect of the project. Y.S., S.L., L.L. performed RNA-seq and some data analysis. W.H. and Y.H. helped with cultures. A.G. M.X., W.Z., and K.M.C. contributed to MEA data collection and analysis. X.S.Y., G.C. and F.R.R contributed to gene expression analysis. H.N.N. and Z.W. generated isogenic iPSC lines. G.C. contributed to Ca^2+^ imaging data collection. A.S. contributed fibroblasts from schizophrenia patients and matched controls. H.S., G-l.M. and Z.G. designed the project and wrote the manuscript.

## Author information

RNA-seq data were deposit at GEO (GSE276169). The authors declare no competing financial interests. Correspondence and requests for materials should be address to Z.G.

## METHODS

### Generation, culture and characterization of patient iPSC lines and isogenic iPSC lines

Skin- punch biopsy samples of three individuals (C3, D2 and D3) were collected from a previously characterized American family, pedigree H^35^. Idiopathic schizophrenic patients (SZ49, SZ59 and SZ69) and normal health subjects (C65, C67 and C75) with matched age, gender and race (Extended Data Table 1), were obtained at Johns Hopkins University School of Medicine. Informed consents were given by each individual participant. All studies followed approved the Stem cell Research Oversight Committee (SCRO), Institutional Review Board (IRB) and animal protocols by Johns Hopkins University School of Medicine and Perelman School of Medicine at the University of Pennsylvania. Mouse embryonic fibroblasts (MEFs) were dissected from E13.5 CF-1 mouse embryos, and were cultured in Dulbecco’s Modified Eagle Medium/Nutrient Mixture F-12 (DMEM/F- 12, Invitrogen) supplemented with 10% fetal bovine serum (FBS, HyClone), 2 mM L-glutamine (Invitrogen), 0.1 mM NEAA (Invitrogen) as previously described^22,34^.

All iPSCs were generated with the EBV-based vectors, and detailed procedures were described before^22,34^. Generally, three vectors: pEP4 EO2S ET2K (Addgene Plasmid 20927), pEP4 EO2S EN2L (Addgene Plasmid 20922), and pEP4 EO2S EM2K (Addgene Plasmid 20923) were transfected in to ∼2x10^6^ human sample fibroblasts by Amaxa Nucleofector (Lonza; program U-023) at a concentration of 2 µg/100 ml electroporation solution. All iPSC colonies were manually picked under the dissect microscope after 3–6 weeks for further expansion and characterization. Colonies with lack of vector integration, was verified by qRT–PCR and excluded as described in previous study^22,34^. For routine culture, iPSCs (passage < 40) were plated on irradiated MEFs in human iPSC medium consisting of DMEM/F12 (Invitrogen), 20% Knockout Serum Replacement (KSR, Invitrogen), 2 mM L-glutamine (Invitrogen), 0.1 mM MEM NEAA (Invitrogen), 0.1 mM ß- mercaptoethanol (Invitrogen), and 10 ng/ml human basic FGF (bFGF, PeproTech). Media were changed daily, and iPSC clones were lifted by collagenase (Invitrogen, 1 mg/ml in DMEM/F12 for 30 min at 37 °C) as described^22,34^. For feeder-free culture of iPSCs, colonies were cultured on Matrigel coated 6-well plate (BD Biosciences) with mTeSR1 media (StemCell Technologies).

C3 and D3 isogenic lines were generated by TALEN in our previous study^22^. TALEN was designed based on a Golden Gate Assembly protocol with modifications to the vector backbone^57^. To generate D2 isogenic line, donor DNA vectors, which were amplified from genomic DNA of a healthy subject, with a loxP-flanked PGK-hygromycin cassette were cloned between 5’ and 3’ homology arms. For targeting, TALENs (4 mg DNA of each plasmid) and linearized donor vectors (10 mg DNA) were electroporated into 1x10^6^ to 2x10^6^ iPS cells, which are pretreated with 5 µM ROCK inhibitor (Y-27632, Cellagentech), using Nucleofector 2b (Lonza; program A-023).

Transfected cells were plated onto a 6-well plate, which is pre-plated with inactivated MEFs. And routine iPSC medium was supplemented with Y-27632 for cell survival. Positive colonies were selected by 10 µg/ml hygromycin B (Invitrogen) after ∼5 days of culture or until small colonies appeared. The selected resistant colonies were then sub-cloned and expanded in 48-well plates. Over 200 clonal lines were screened per individual. To remove loxP-flanked PGK-hygromycin cassette, the electroporation of a Cre recombinase expression vector (4 mg DNA) was performed. Specific integration, correct genetic editing and efficient removal of PGK-hygromycin cassette were confirmed by Sanger sequencing at each stage.

Karyotyping analysis by standard G-banding technique was carried out by Cell Line Genetics and the results were interpreted by clinical laboratory specialists of Cell Line Genetics. Genotyping of all DISC1 lines except D2R was performed and analyzed in our previous study^22^. Genomic DNA of D2R line was extracted by DNeasy Blood & Tissue Kit (Qiagen) following the manufacturer’s recommended protocol. A pair of primers was designed and used to amplify the region around the 4-bp deletion. PCR products were cloned by TA cloning and sequenced.

### Differentiation of iPSCs into forebrain cortical neurons and GABAergic neurons

The protocol of differentiation of iPSCs into forebrain-specific neural progenitors and cortical neurons was described previously^22^. The purmorphamine (Pur, StemGent), a smoothened activator was used to pattern medial ganglionic eminence (MGE) progenitors referring to the previous protocol^58^.

Specifically, to generate MGE progenitors and GABAergic neurons, iPSC colonies were lifted from the feeder layer with 1 mg/ml collagenase treatment for 30 min and suspended in embryoid body (EB) medium, consisting of FGF-2 free iPSC medium supplemented with 2 µM dorsomorphin and 2 µM A-83, in non-treated polystyrene plates for 4 days with a daily medium change. After 4 days, EB medium was replaced by neural induction medium (NIM) consisting of half DMEM/F12 and half Neurobasal medium, N2 supplement, NEAA and high concentration of Pur (1.5 µM). The floating EBs were then transferred to Matirgel-coated 6-well plates at day 7 to form neural tube-like rosettes. The attached rosettes were kept for 15 days with NIM half-change every other day. On day 22, the rosettes were picked mechanically, and transferred to low attachment plates (Corning) to form neurospheres in NIM medium for 3 days. The suspended neurospheres were dissociated with Accutase at 37 °C for 10 min and place onto Matirgel-coated 6-well plates. The monolayer progenitors will be prorogated for 2-3 passages cultured in NIM media containing low concentration of Pur (0.5 µM) before neuronal differentiation. The isolated MGE progenitors with Accutase at 37 °C for 5 min were placed onto poly-D-lysine/laminin-coated coverslips in the neuronal culture medium, consisting of Neurobasal medium supplemented with 2 mM L-glutamine, B27, 10 ng/ml BDNF (PeproTech), 10 ng/ml GDNF (PeproTech), 200 ng/ml Ascorbic Acid (AA, Sigma) and 1 µM cAMP (Sigma). The compound E (0.2 µM, EMD Biosciences) was applied to synchronize post-mitotic neurons for 3 days. Half of the medium was replaced once a week during continuous culturing. For immune staining and electrophysiological recordings, neural progenitors were plated on a confluent layer of rodent astrocytes as previously described^22,59^. These cultures exhibited similar neuronal densities and parallel cultures were used for examination of different iPSC lines.

### Immunocytochemistry analyses

All samples were fixed with 4% paraformaldehyde (Sigma) for 15 min at room temperature and washed 3 times with PBS. Cells were permeabilized with 0.1% Triton X-100 (Sigma) and blocked in 10% donkey serum for 60 min as previously described^22^. Samples were then incubated with primary antibodies (Extended Data Table 1b) diluted in PBS containing 0.1%Triton X-100 and 10% donkey serum at 4 °C overnight, followed by incubation with secondary antibodies diluted (1:500-1:1000) in 10% donkey serum for 2 h at room temperature.

The samples were visualized by Zeiss LSM710 confocal microscope, Zeiss LSM800 confocal microscope or Zeiss Axiovert 200M microscope. Images were acquired with identical acquisition settings for parallel cultures, and the taken images were analyzed with ImageJ (NIH). For analysis of synaptic bouton density, total SV2, SYN1, GAD65 and vGAT puncta in a given image were counted by ImageJ Analyze Particles, and the total dendritic length were measured by ImageJ plugin NeurphologyJ^60^. The synaptic density was determined by D (D = total SV2, SYN1, GAD65 or vGAT puncta per 100 mm total dendritic length). For quantification of total KCC2 expression, the mean KCC2 intensity was calculated, and was divided by numbers of neurons per image.

### Electrophysiology

All whole-cell patch-clamp recordings were conducted at room temperature and were performed on iPSC-derived neurons using Multiclamp 700A patch-clampamplifier (Molecular Devices, Palo Alto, CA) as previously described^22^. Briefly, neurons grown on coverslips were visualized with a 40X air objective on an inverted Zeiss microscope (Zeiss Axiovert 200M microscope), and the recording chamber was constantly perfused with a bath solution consisting of 128 mM NaCl, 30 mM glucose, 25 mM HEPES, 5 mM KCl, 2 mM CaCl_2_, and 1 mM MgCl_2_ (pH 7.3; 315–325 Osm). Patch pipettes were pulled from borosilicate glass (BF150-86-10, Sutter Instrument) with a resistance of 3-5 MΩ and filled with an internal solution consisting of 135 mM CsCl, 10 mM Trisphosphocreatine, 10 mM HEPES, 2.5 mM EGTA, 4 mM MgATP, 0.5 mM Na_2_GTP and 5 mM QX-314 (pH 7.3 with CsOH; ∼305 Osm; for IPSCs, evoked IPSCs and r-aminobutyric acid (GABA)-current measurements) and 135 mM CsGluconate, 10 mM Trisphosphocreatine, 10 mM HEPES, 2.5 mM EGTA, 4 mM MgATP, 0.5 mM Na_2_GTP and 5 mM QX-314 (pH 7.3 with CsOH; ∼305 Osm; for EPSCs measurements holding at -70mV and for IPSCs measurement holding at + 20 mV). The series resistance was typically 10–30 MΩ. The 10 µM NBQX (Tocris Bioscience) and 50 µM 2-amino-5-phosphonopentanoic acid (D-AP5, Tocris Bioscience) were present in the extracellular solutions to isolate spontaneous IPSCs, evoked IPSCs and GABA-current from focal application. The 20 µM bicuculline (Bic, Tocris Bioscience) were applied for spontaneous EPSC recording. Drugs were applied through a gravity driven drug delivery system (VC-6, Warner Hamden, CT). Spontaneous IPSCs and EPSCs were recorded for 4 min (2 min after break-in to block sodium current by QX-314) at the holding potential, sampled at 10 kHz and filtered at 1 kHz. Evoked postsynaptic currents were induced by brief (1 ms) unipolar current pulses of various amplitudes (0.1–0.9 mA, delta = 0.1 mA). The current pulses were delivered with a stimulating electrode (CBAEC75, FHC), which was positioned about 100–150 μm from the cell soma and connected to an external stimulator (A365, WPI). To measure pair-pulse ratio (PPR), the mean Peak2/mean Peak1 was calculated at a stimulus interval of 100 ms. GABA-currents were induced by a focal transient application of the following 100 µM GABA. An injection pipette (BF150-110-10, Sutter Instrument), with resistance of 1–2 MΩ when filled with extracellular solution was positioned about 20 μm from the cell soma and applied focally by in-line pressure application (10 p.s.i., Pressure System IIe, Toohey Company). Data were acquired using pClamp 9 software (Molecular Devices, Palo Alto, CA), and spontaneous synaptic events were analyzed using MiniAnalysis software (Synaptosoft, Decatur, GA).

### Calcium Image

iPSC-derived neurons were incubated in 2.5 μM Fura-2-acetoxymethyl ester (Fura-2-AM, Invitrogen) for 50 min at 37°C, and washed for 20 min in bath solution. The Fura-2-AM labeled neurons on coverslips were transferred to a perfusion chamber mounted on a Nikon TE-2000-S inverted microscope with a 20X objective and imaged with a 340/380 nm transmittance filter set (Chroma Technology). SimplePCI (HCImage, Hamamatsu) was used to measure the ratio of 340/380 fluorescence signals in neuronal soma. Sister coverslips from 3 independent cultures were taken for imaging at 4-week and 6-week time points. All imaging was performed in the presence of DNQX (10 μM). The threshold of a significant calcium response was set as 10 times of baseline standard deviation (STDV).

### Micro Electrode Array (MEA)

To examine neuronal network, C3 NPC, C3 MGE, D3 NPC, and D3 MGE were co-cultured as four different combinations (C3 NPC/ C3 MGE, C3 NPC/ D3 MGE, D3 NPC/ C3 MGE and D3 NPC / D3 MGE) at ratio of 80 to 20. The mixed progenitors were plated into a 12-well MEA plate (Axion Biosystems). Each well contained an 8 × 8 grid of 30 nm circular nanoporous platinum electrodes embedded in the cell culture substrate. Wells were then coated in PDL (20 µg/ml) and laminin (10 µg/ml) and cells were plated at 400,000 cells/well. MEA plates were incubated at 37°C and 5% CO_2_, and sterile water was added into extra-well space to prevent media evaporation. Neurons were cultured in Brainphys media (STEMCELL Technologies) containing SM1 neuronal supplement (STEMCELL Technologies), 2 mM L-glutamine, B27, 10 ng/ml BDNF, 10 ng/ml GDNF200 ng/ml AA and 1 µM cAMP, treated by Compound E at day 1 to synchronize neurons, and fed every week. Extracellular recordings of spontaneous action potentials were performed in Brainphys medium at 37°C using a Maestro MEA system and AxIS software (Axion Biosystems). Data were sampled at rate of 12.5 kHz with a hardware frequency bandwidth of 200- 3000 Hz. For unsupervised spike detection, the threshold was set to 6 times STDV of the base line on each electrode using Neuroexplore. The algorithm of individual bursts on each electrode were defined as Extended Data Fig. 6b. The custom Python codes were used to generate heat-map and to analyze network synchrony. Five-minute recordings were used to calculate mean spike rate and network synchrony for the wells.

### RNA-seq analysis

Total RNA was purified from 6-week-old GABAergic neurons using Rneasy kit (Qiagen). Dnase I on-column digestion was performed to avoid genomic DNA contamination.

Sequencing libraries were prepared using NEBNext Ultra RNA Library Prep kit for Illumina (E7530L) following manufacturer’s protocol. Briefly, total RNA was poly-A tail selected and then heat fragmented. The fragmented RNA was reverse transcribed, and the second strand was synthesized to make double stranded DNA. After end repair and 3’ adenylation, adapters for multiplexing were ligated to the end of double stranded DNA fragments. The ligation products were amplified and purified to generate illumina compatible libraries. Sequencing was performed with 75bp single end sequenced by illumina NextSeq 550. The raw reads were mapped to the human genome build hg38 using tophat^61^. The differential gene expression and downstream analyses were performed using DESeq2^62^ and custom R scripts. Significantly dysregulated genes were defined as p value < 0.01 and fold change >1.8. The RNA-seq statistics and lists of differentially expressed genes are summarized in Extended Data Table 2. Pathway signature enrichment analysis and disease enriched categories were performed using online source DAVID v6.8 (https://david.ncifcrf.gov/).

### Data collection and statistics

All experiments were replicated at least three times using iPSC lines indicated in Extended Data Table 1, and data were acquired from parallel cultures. The sample size and description of the sample collection are described in each figure legend. Statistical tests used for comparison are reported in all figure legends. For data sets where parametric tests were used for analysis, the normal distribution of data was confirmed using the Kolmogorov– Smirnov test.

## Supplementary Information

**Extended Data Figure 1. Basic characterization of the pedigree H iPSC lines and forebrain neurons**. **a**, A schematic diagram of the pedigree H for iPSC generation and isogenic iPSC lines in the current study. The symbol + indicates one copy of the 4-bp deletion in the *DISC1* gene; the symbol - indicates a lack of the 4-bp deletion in the ***DISC1*** gene. **b**, Sample confocal images of immunostaining for pluripotency markers (scale bar, 50 µm) and sample image of karyotyping for D2R line. The same analyses for other lines used in the current study were previously reported^22^. **c-f**, Differentiation of iPSC lines into forebrain cortical neural progenitors. Shown are sample confocal images of immunostaining for nestin and forebrain progenitor markers, PAX6 (**c**) and OTX2 (**e**) (scale bar, 40 µm) and quantification of PAX6^+^ (**d**) and OTX2^+^ (**f**) progenitors among different iPSC lines. Values represent mean ± s.e.m. (n = 4 to 7 cultures). **g-h**, Presence of GABAgeric neurons in forebrain neuronal cultures. Shown are sample confocal images of immunostaining for GAD67 of human forebrain neurons at 4 weeks after neuronal differentiation (**g**; scale bar, 40 µm) and quantification of GAD67^+^ neurons among different iPSC lines. Values represent mean ± s.e.m. (n = 5 to 17 cultures).

**Extended Data Figure 2. Defects of GABAergic synaptic transmission in *DISC1* mutant forebrain neurons. a-c,** Sample recording traces and cumulative distribution plots of spontaneous IPSCs intervals and amplitudes of isogenic pairs of forebrain neurons at 4 weeks after neuronal differentiation (n = 6 to 14 neurons for each condition; Kolmogorov–Smirnov test). **d-f,** Cumulative distribution plots of spontaneous IPSCs intervals and amplitudes of 6-weeks old forebrain neurons with isogenic paired comparison (n = 5 to 7 neurons for each condition; Kolmogorov–Smirnov test).

**Extended Data Figure 3. Ganglionic eminence-specific GABAergic neural differentiation of iPSC lines. a,** A schematic diagram of the differentiation procedure**. b-e,** Sample images of immunostaining for MGE progenitor markers, FOXG1 (**b**) and NKX2.1 (**d**) (scale bar, 40 µm) and quantifications of FOXG1^+^ (**c**) and NKX2.1^+^ (**e**) neural progenitors among different iPSC lines. Values represent mean ± s.e.m. (n = 5-8 cultures).

**Extended Data Figure 4. Decreased density of vGAT^+^ puncta and defects of synaptic transmission in *DISC1* mutant GABAergic neurons. a,** Sample confocal images of vGAT and MAP2 immunostaining of 6-weeks old GABAergic neurons (scale bar, 5 µm). **b**, Quantification of vGAT^+^ puncta density. Values represent mean ± s.e.m. (n = 10-16 cultures; ***p < 0.001; Student’s t-test). **c-e**, Cumulative distribution plots of spontaneous IPSCs intervals and amplitudes of 6-weeks old GABAergic neurons with isogenic paired comparison (n = 6 to 29 neurons for each condition; Kolmogorov–Smirnov test).

**Extended Data Figure 5. *DISC1* mutant GABAergic neurons exhibit dysregulated synaptic transmission at 8 weeks after neuronal differentiation. a-c,** Sample recording traces and cumulative distribution plots of spontaneous IPSCs intervals and amplitudes of isogenic paired GABAergic neurons at 8 weeks after neuronal differentiation (n = 10 to 22 neurons for each condition; Kolmogorov–Smirnov test)**. d,** Sample recording traces of evoked IPSCs of 8-weeks old GABAergic neuron with isogenic paired comparison and quantification. Values represent mean ± s.e.m. (n = 4 to 11 neurons for each condition;***p < 0.001; **p < 0.01, Student’s t-test). **e,** Sample recording traces and summary bar graph showing pair-pulse ratio. Values represent mean ± s.e.m. (n = 5 to 13 neurons for each condition; ** p < 0.01; * p < 0.05, Student’s t-test)**. f,** Sample recording current tracers of control and *DISC1* mutant GABAergic neurons in response to focal application of GABA (100 µM) and quantification of GABA induced currents. Values represent mean ± s.e.m. (n = 4 to 8 neurons for each condition; *** p < 0.001; * p < 0.05, Student’s t-test).

**Extended Data Figure 6. *DISC1* mutant glutamatergic and GABAergic neuron co-culture displays abnormal neuronal network activity and synchronization. a,** Sample bright-field image of human glutamatergic and GABAergic neuron co-culture at 6 weeks after neuronal differentiation (scale bar, 200 µm). **b,** Schematic diagram of the burst definition on a single electrode. **c,** Sample recording traces of spikes and bursts on a single electrode. Blue boxes present burst duration and raster plots of spike activity are also shown below. **d-f,** Cumulative distribution plots of spike frequency (**d**), burst duration (**e**) and number of spikes in burst (**f**) under four different co-culture combinations at 6 weeks after neuronal differentiation (n = 3 cultures; **p < 0.01, ***p < 0.001; Kolmogorov–Smirnov test;).

**Extended Data Figure 7. Functional switch of GABA responses in human neurons at 6 weeks after neuronal differentiation. a,** Sample images showing 6-weeks old human neurons loaded with Fura-2 (scale bar, 40 µm). **b,** Sample traces showing somatic Ca^2+^ responses after stimulation with 100 μM GABA and 90 mM KCl in C3 forebrain neurons at 4 and 6 weeks. **c-d,** Bar graphs showing the percentages of neurons showing GABA-evoked Ca^2+^ increase (**c**) and the amplitude of GABA-evoked Ca^2+^ responses (**d**). Values represent mean ± s.e.m. (n = 3 cultures) **e,** Sample confocal images showing reduced KCC2 immunostaining (confocal) in 6-weeks old neurons compared to 4-weeks (scale bar, 40 µm). **f,** Quantification of KCC2 immunofluorescence intensity. Values represent mean ± s.e.m. (n = 4 to 6 cultures).

**Extended Data Figure 8. Basic characterization of iPSC lines derived from idiopathic schizophrenia patients and age, gender and race matched control subjects. a,** Sample confocal images of pluripotency markers (scale bar, 50 µm). **b**, sample image of karyotyping for both control and idiopathic schizophrenia patient iPSC lines.

**Extended Data Figure 9. Forebrain neural differentiation of idiopathic schizophrenia patient and matched control iPSC lines. a-b,** Sample confocal images of immunostaining of nestin and forebrain progenitor makers, Pax6 and Otx2 (**a**; scale bar, 40 µm) and quantification (**b**). Values represent mean ± s.e.m. (n = 4 cultures). **c-d,** Sample images of immunostaining of forebrain neuronal markers, Tbr1, Ctip2 and Brn2 (**c**, scale bar, 40 µm) and quantification (**d**). Values represent mean ± s.e.m. (n = 5 cultures).

**Extended Data Figure 10. Characterization of neuronal subtypes derived from idiopathic schizophrenia patient and matched control iPSC lines. a-b,** Expression of glutamatergic neuron maker vGlut1. Shown are sample confocal images of immunostaining of vGlut1 and DCX (**a**; scale bar, 40 µm) and quantification (**b**). Values represent mean ± s.e.m. (n = 5 cultures), **c, d,** Expression of GABAergic neuron marker GAD67. Same as in **a-b**, except that GAD67 was examined.

**Extended Data Figure 11. Dysregulation of both excitatory and inhibitory synaptic transmission in forebrain neurons derived from idiopathic schizophrenia patient iPSCs compared to matched controls. a-c,** Defects in glutamatergic synaptic transmission in idiopathic schizophrenia patient iPSC-derived forebrain neurons in comparison to control neurons. Shown are sample whole-cell voltage-clamp recording traces of excitatory postsynaptic synaptic currents (EPSCs) of 6-weeks old forebrain neurons (**a**) and cumulative distribution plots of spontaneous EPSCs intervals (**b**) and amplitudes (**c**). Also shown are quantifications of mean frequencies and amplitudes of spontaneous EPSCs. Values represent mean ± s.e.m (n = 9 to 14 neurons for each condition; **p < 0.01 and n.s. denotes non-significance; Student’s t-test). **d-f**, Defects in GABAergic synaptic transmission in idiopathic schizophrenia patient iPSC-derived forebrain neurons in comparison to control neurons. Shown are sample whole-cell voltage-clamp recording traces of spontaneous inhibitory post synaptic currents (IPSCs) of 6-weeks old forebrain neurons (**d**; V_m_ = 20 mV) and cumulative distribution plots of spontaneous IPSCs intervals (**e**) and amplitudes (**f**). Also shown are quantification of mean frequencies and amplitudes of spontaneous IPSCs. Values represent mean ± s.e.m. (n = 7 to 16 neurons for each condition; **p < 0.01, ***p < 0.001; Student’s t-test).

**Extended Data Figure 12. Differentiation of idiopathic schizophrenia patient and matched control iPSC lines into GABAergic neurons. a-b,** Characterization of MGE progenitors derived from different iPSC lines. Shown are sample confocal images of immunostaining of nestin and MGE progenitor maker NKX2.1 and progenitor marker Sox2 (**a**; scale bar, 40 µm) and quantification (**b**). Values represent mean ± s.e.m. (n = 4 to 5 cultures). **c-d**, Characterization of GABAergic neurons derived from different iPSC lines. Shown are sample confocal images of immunostaining of GAD67 and DCX (**c;** scale bar, 40 µm) and quantification (**d**). Values represent mean ± s.e.m. (n = 4 to 7 cultures).

**Extended Data Figure 13. Dysregulation of neuronal transcriptome in *DISC1* mutant GABAergic neurons. a,** MA plots showing differentially expressed genes of 6-weeks old GABAergic neurons (C3 and D3 pairs; n = 3 to 4 samples for each iPSC line, p value < 0.01 and fold change >1.8). **b**, Venn diagrams comparing the overlap in differential gene expression between C3 and D3 pair (significantly overlapping was tested by fisher test; up regulation p value = 1.8e- 123, down regulation p value = 3.9e-97). **c**, ReViGo scatter plots of GO analyses for GABAergic neurons from *DISC1* mutant and isogenic control lines. Bubble colors represent p-values and bubble sizes indicate the relative frequency of the GO terms. **d-e**, Enrichment analysis by disease categories. Bubble plots showing top enriched disease categories for differentially expressed genes in GABAergic neurons of *DISC1* mutant and isogenic control lines. Each dot represents one dataset, and a table summarizes statistics for meta-comparison.

**Extended Data Figure 14. Transcriptome dysregulation of idiopathic schizophrenia patient iPSC-derived GABAergic neurons. a,** ReViGo scatter plots of GO analyses for GABAergic neurons from idiopathic schizophrenia patient and matched control cohorts. Bubble colors represent p-values and bubble sizes indicate the relative frequency of the GO terms. **b-c**, Enrichment analysis by disease categories and in different mental disorders and nervous system diseases. Similar as in Extended Data Figure 13d-e. Bubble plots showing top enriched disease categories for differentially expressed genes in GABAergic neurons of idiopathic schizophrenia patient and matched control cohorts. Each dot represents one dataset, and a table summarizes statistics for meta-comparison.

**Extended Data Table 1. Summary of reagents in the current study. a,** Characterization of all iPSC lines used in the current study. **b**, List for antibodies used in the current study.

**Extended Data Table 2. Summary of RNA-seq analysis. a,** Summary of sequencing information. **b**, List of differentially expressed genes between isogenic pair C3M and C3; **c**, List of differentially expressed genes between isogenic pair D3 and DR3. **d**, List of differentially expressed genes between idiopathic schizophrenia patient iPSC-derived GABAergic neurons and those derived from matched controls.

