## Supplementary figures and images for "GABAergic neuron dysregulation in a human neurodevelopmental model for major psychiatric disorders"

### Supplemental Figure 1

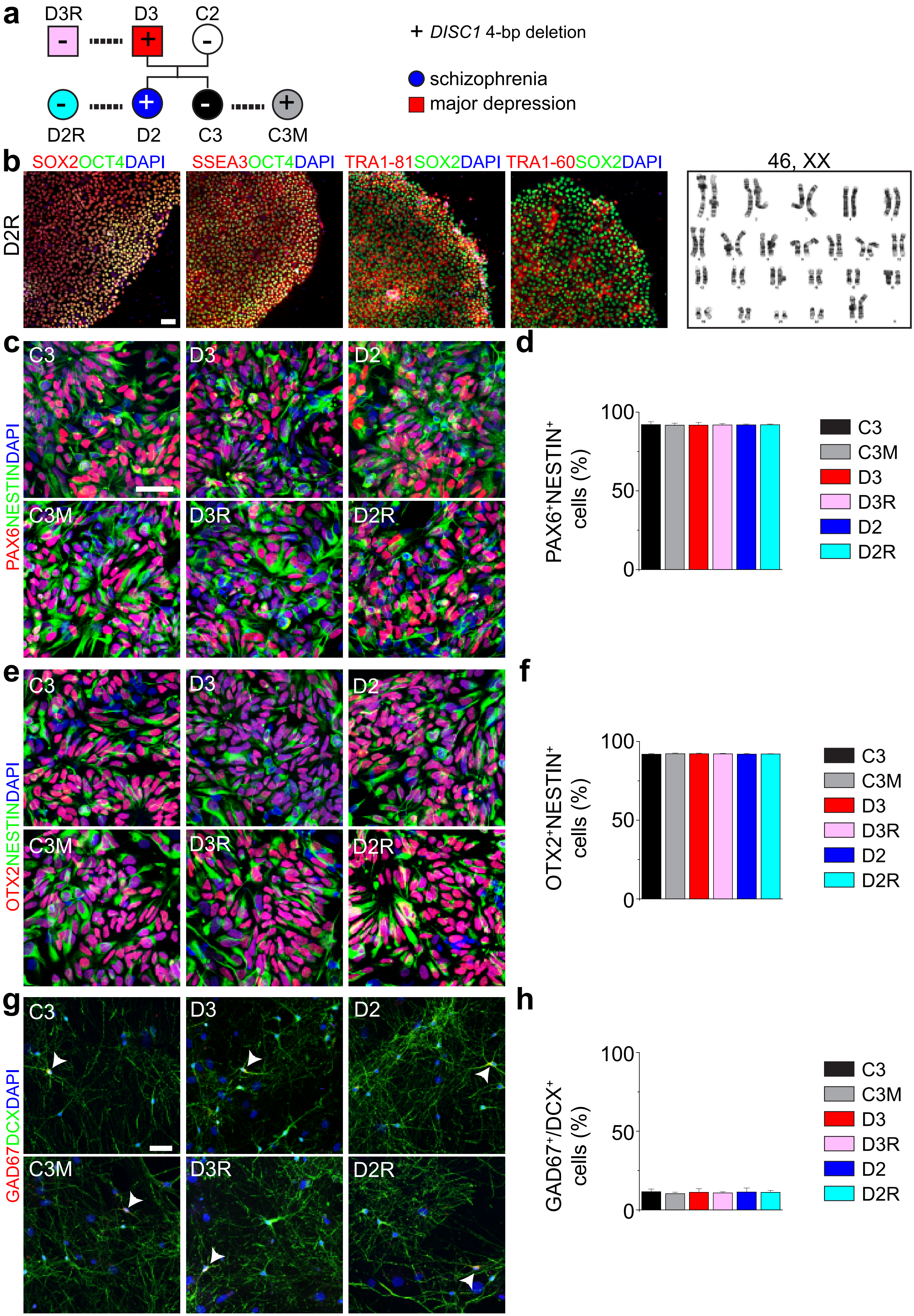


Supplementary Figure 1 (Guo et al.,)

### Supplemental Figure 2

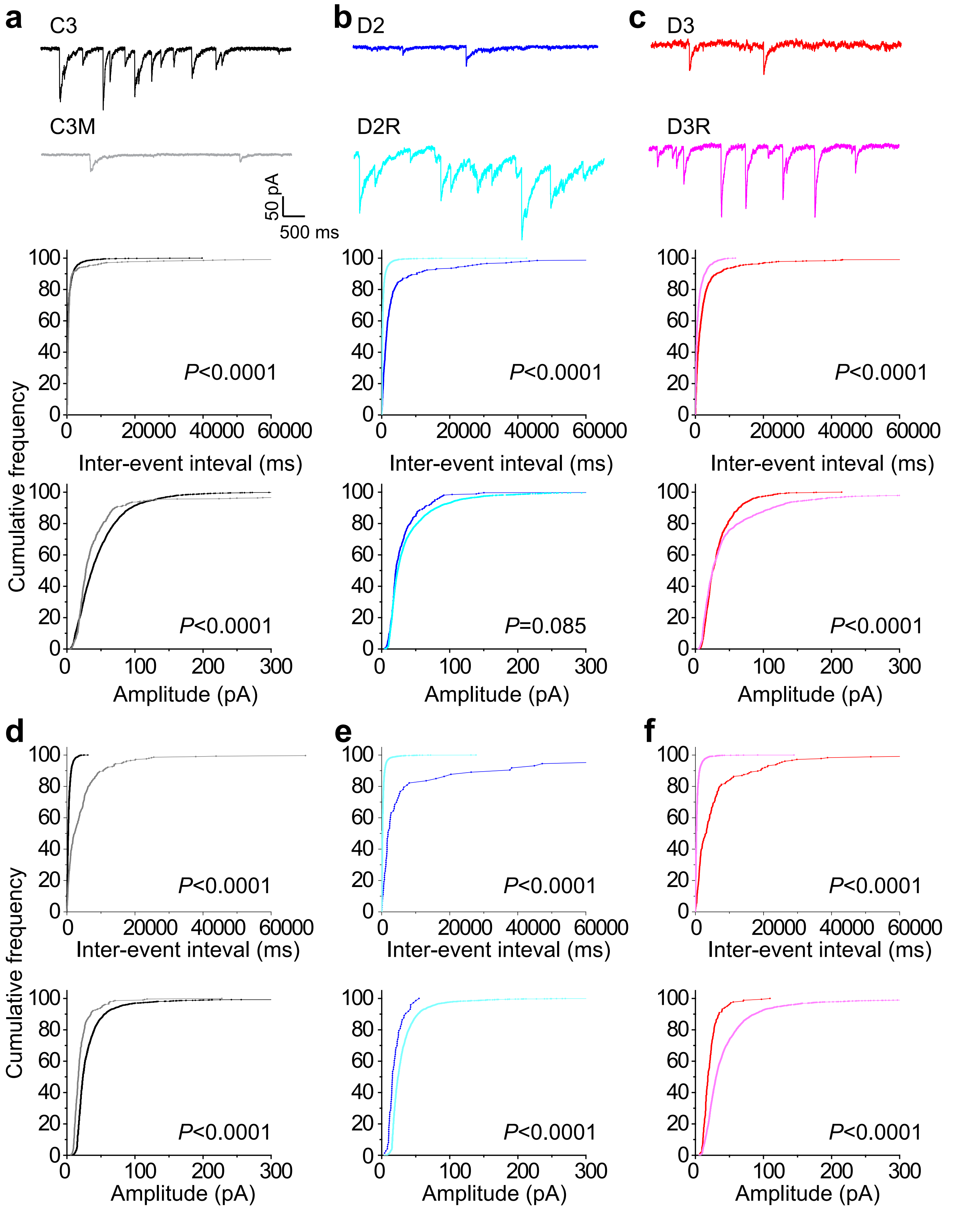


Supplementary Figure 2 (Guo et al.)

### Supplemental Figure 3

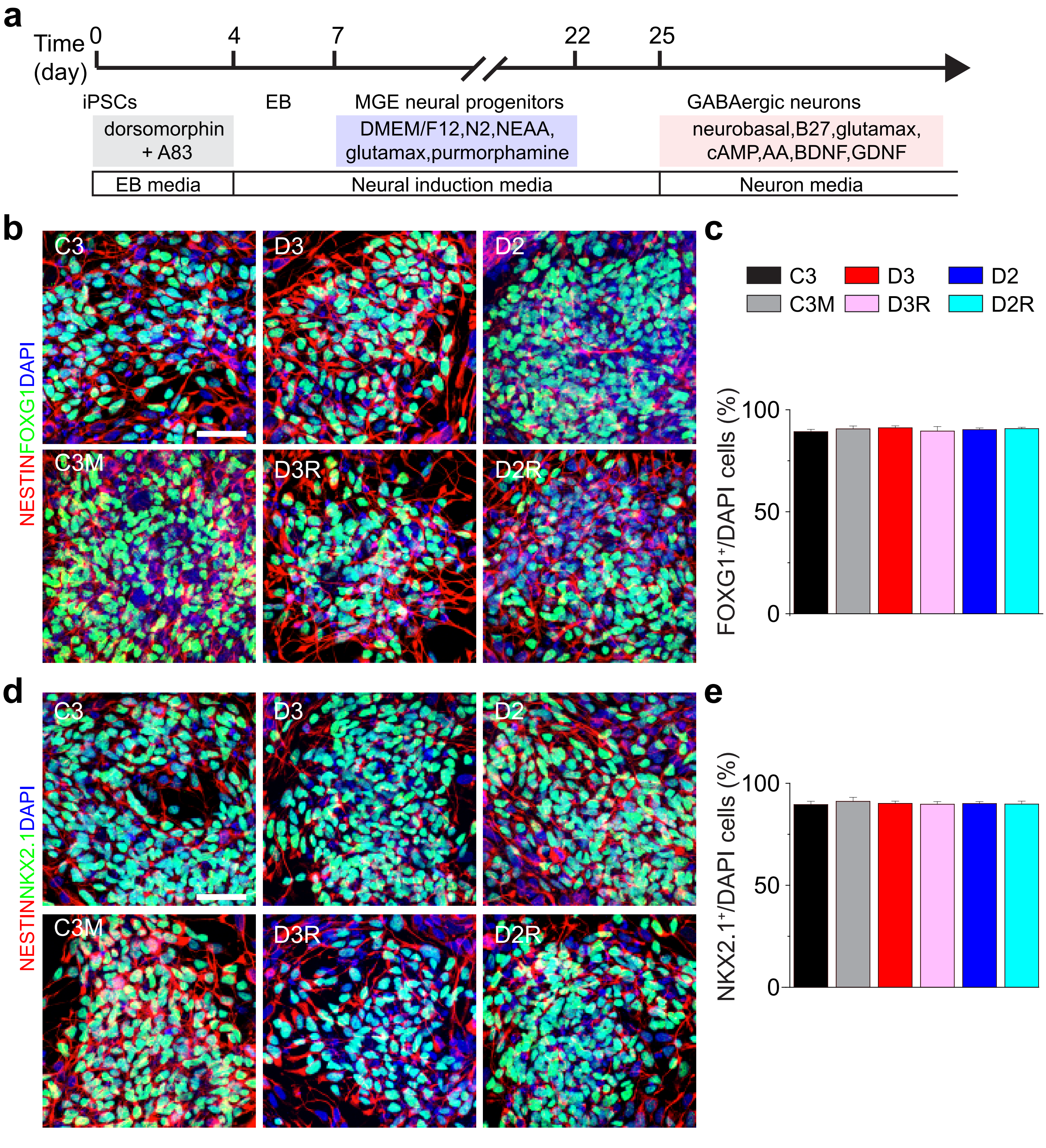


Supplementary Figure 3 (Guo et al.,)

### Supplemental Figure 4

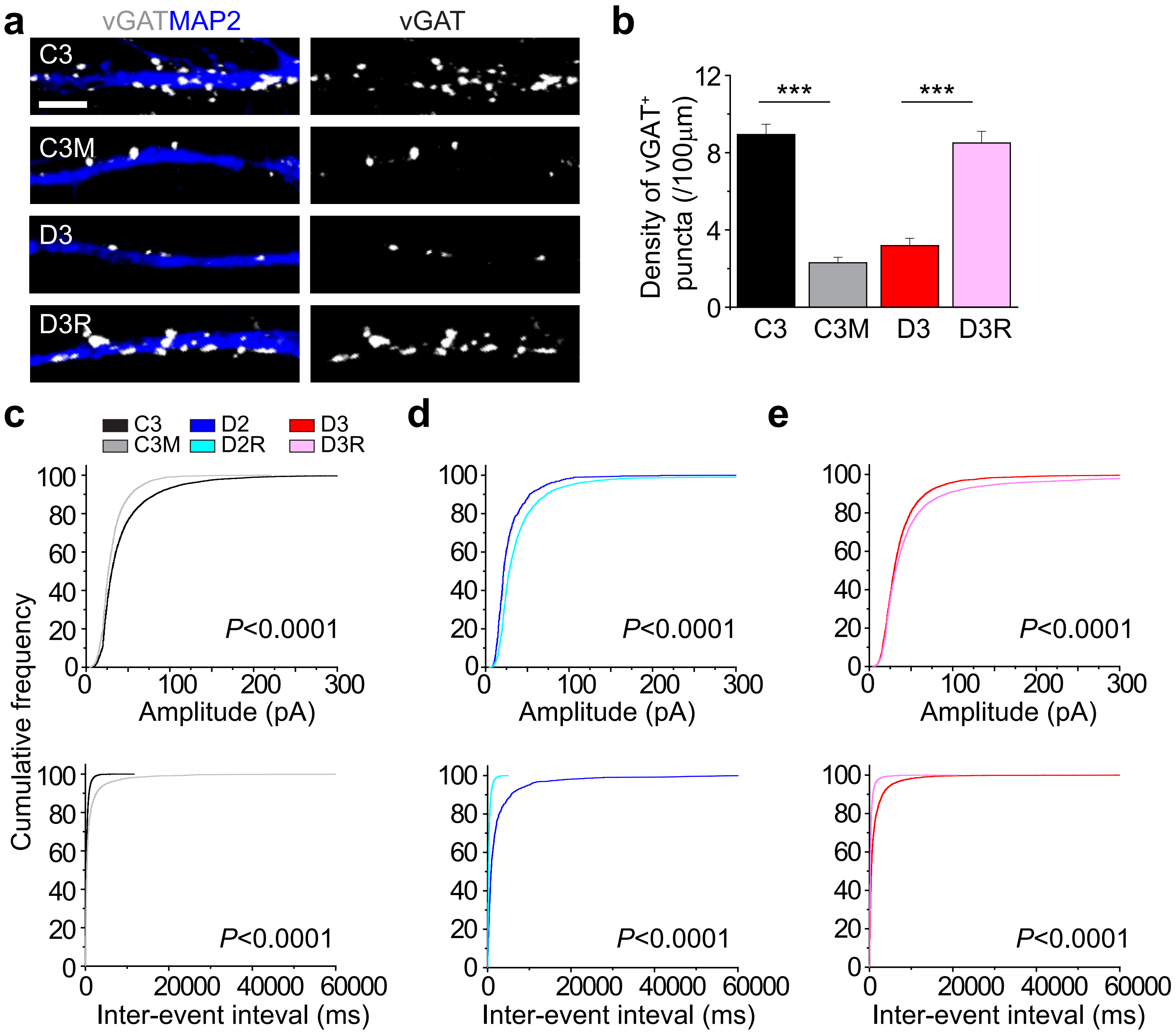


Supplementary Figure 4 (Guo et al.,)

### Supplemental Figure 5

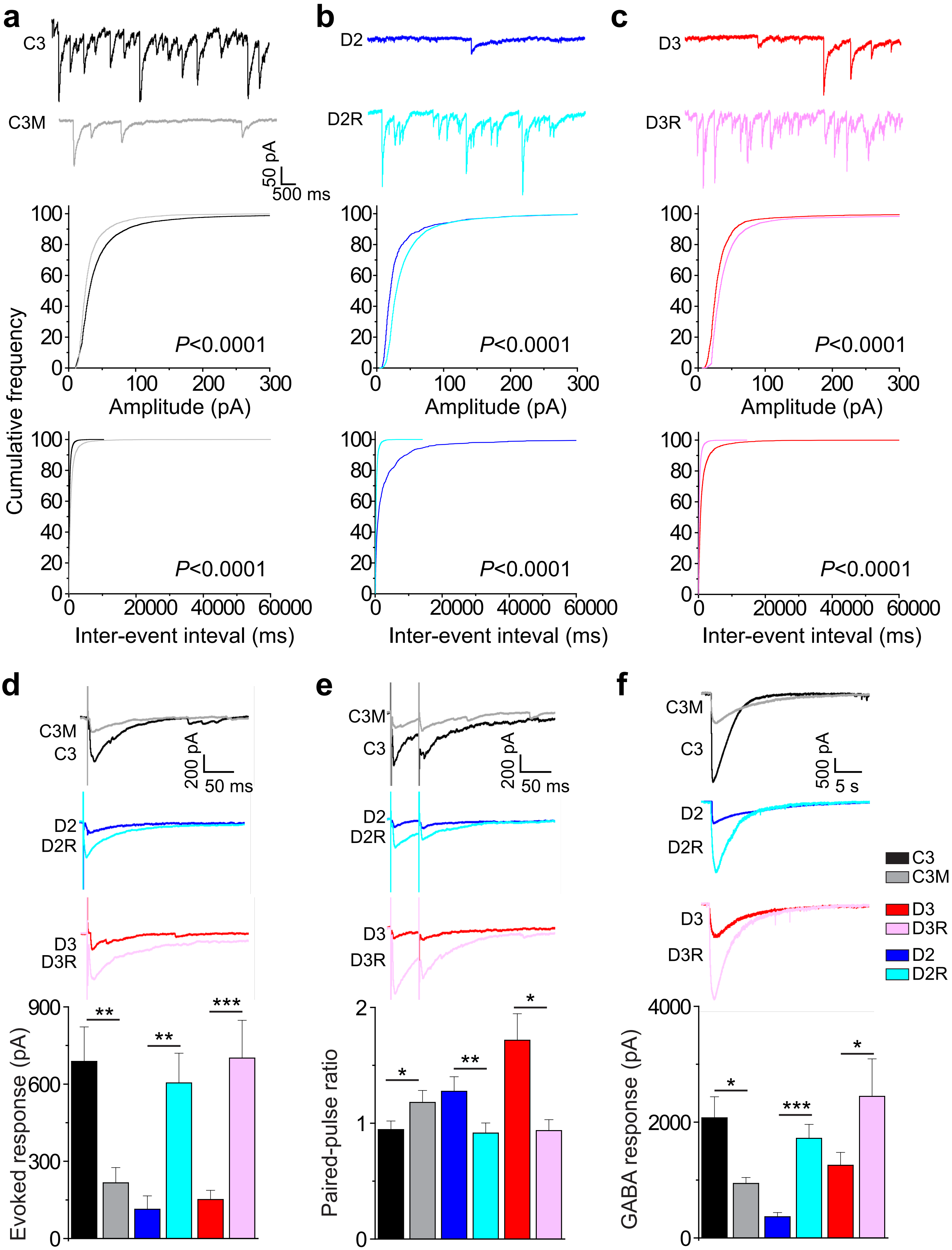


Supplementary Figure 5 (Guo et al.,)

### Supplemental Figure 6

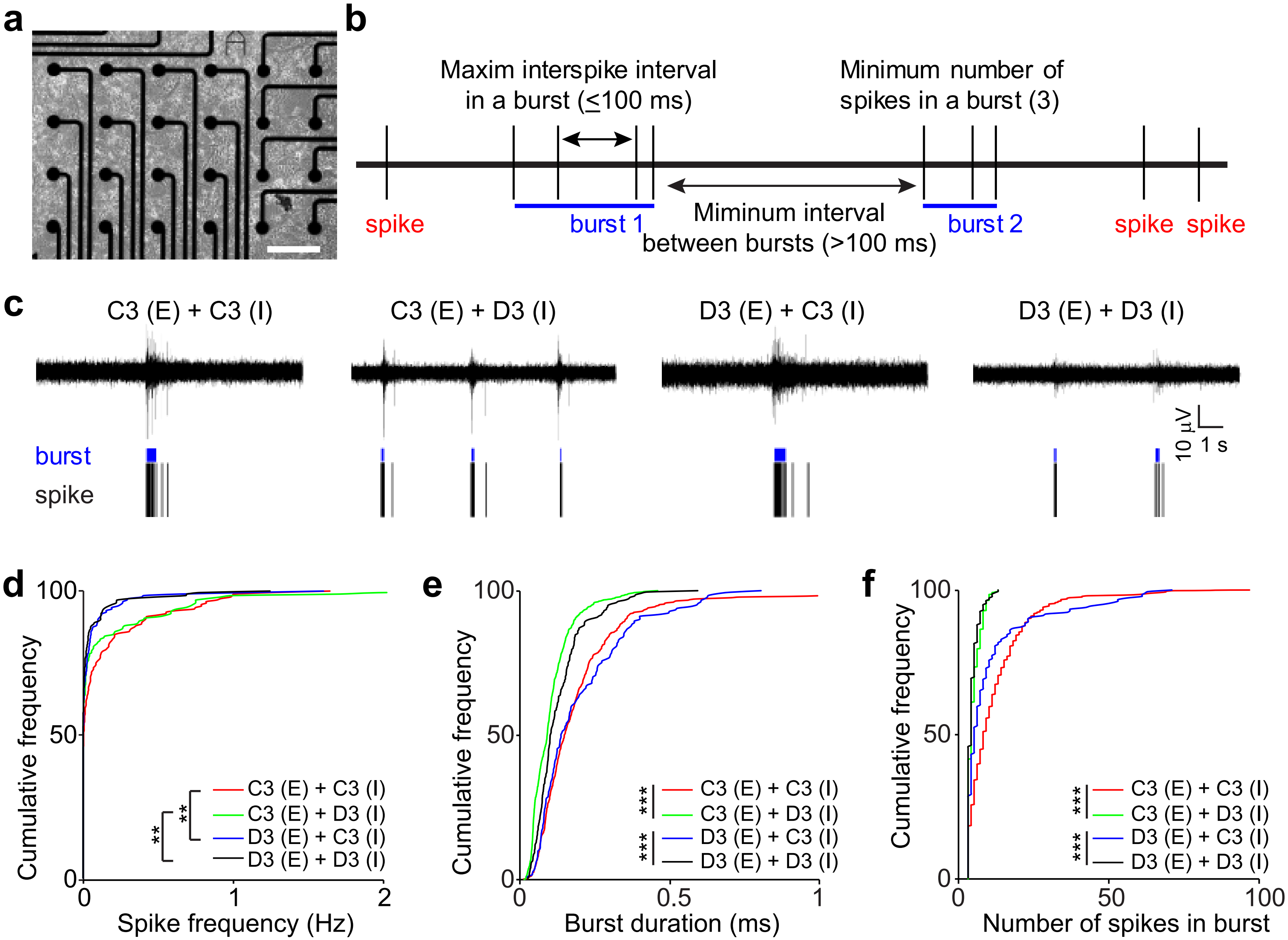


Supplementary Figure 6 (Guo et al.,)

### Supplemental Figure 7

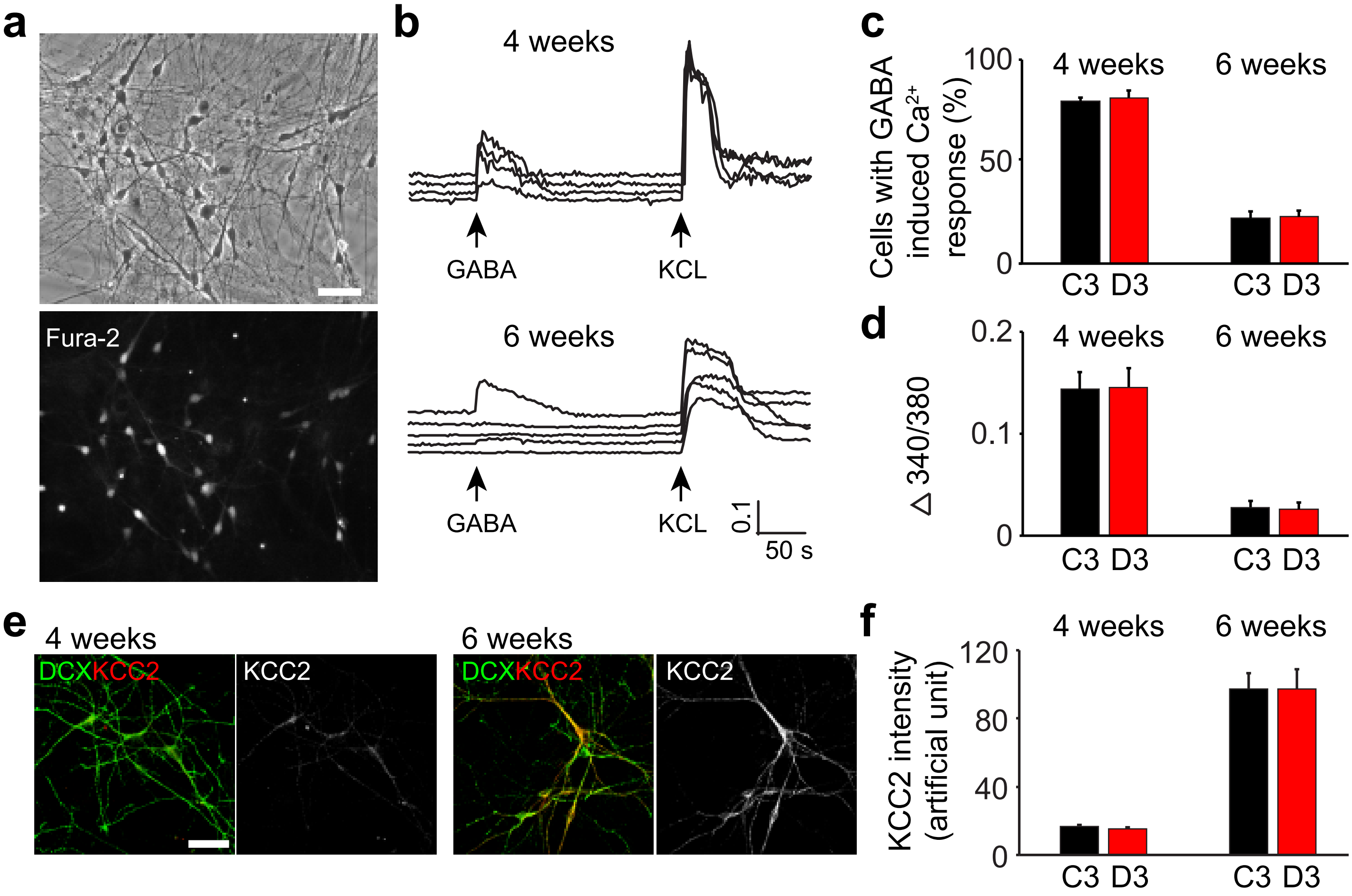


Supplementary Figure 7 (Guo et al.,)

### Supplemental Figure 8

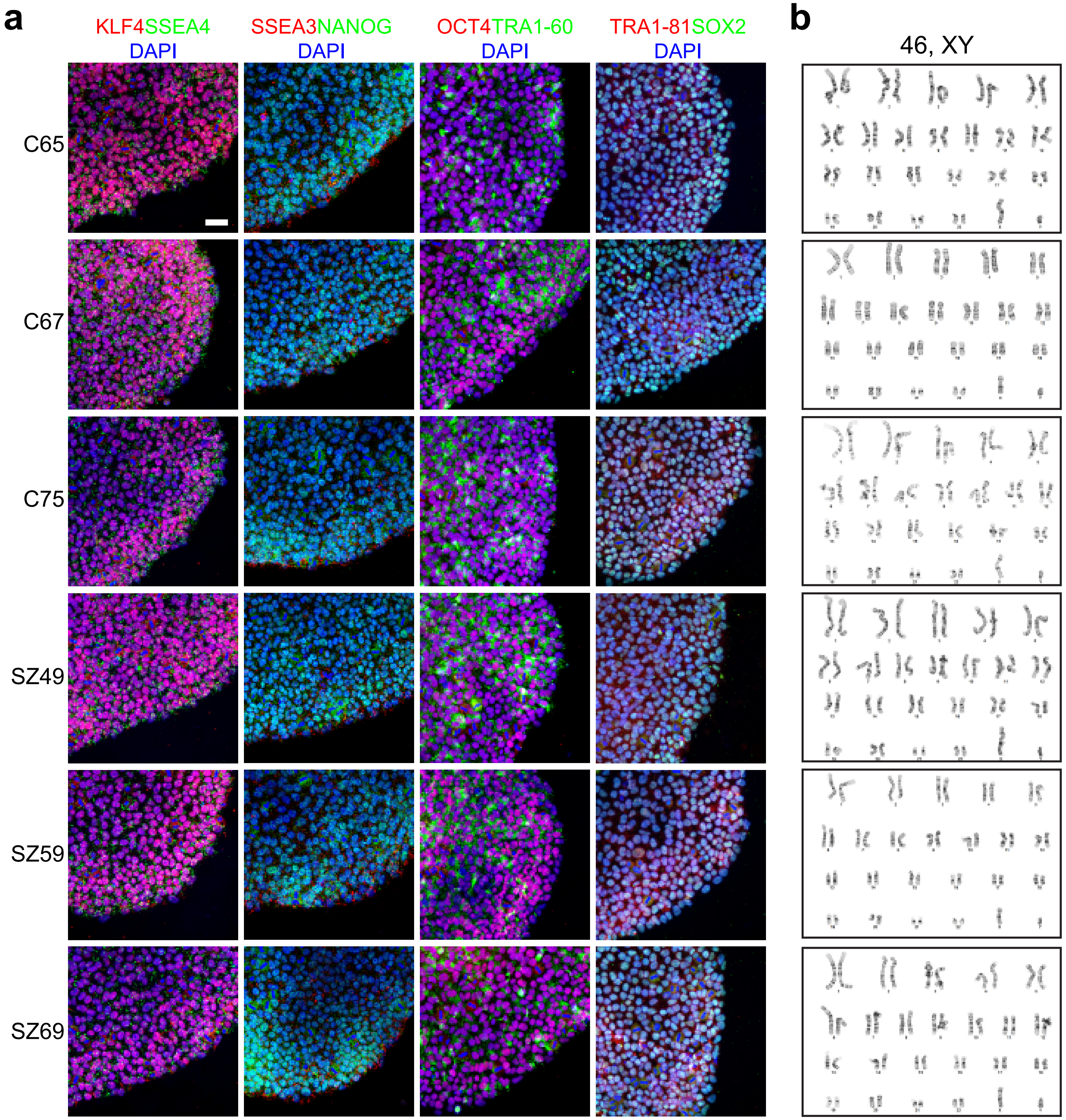


Supplementary Figure 8 (Guo et al.,)

### Supplemental Figure 9

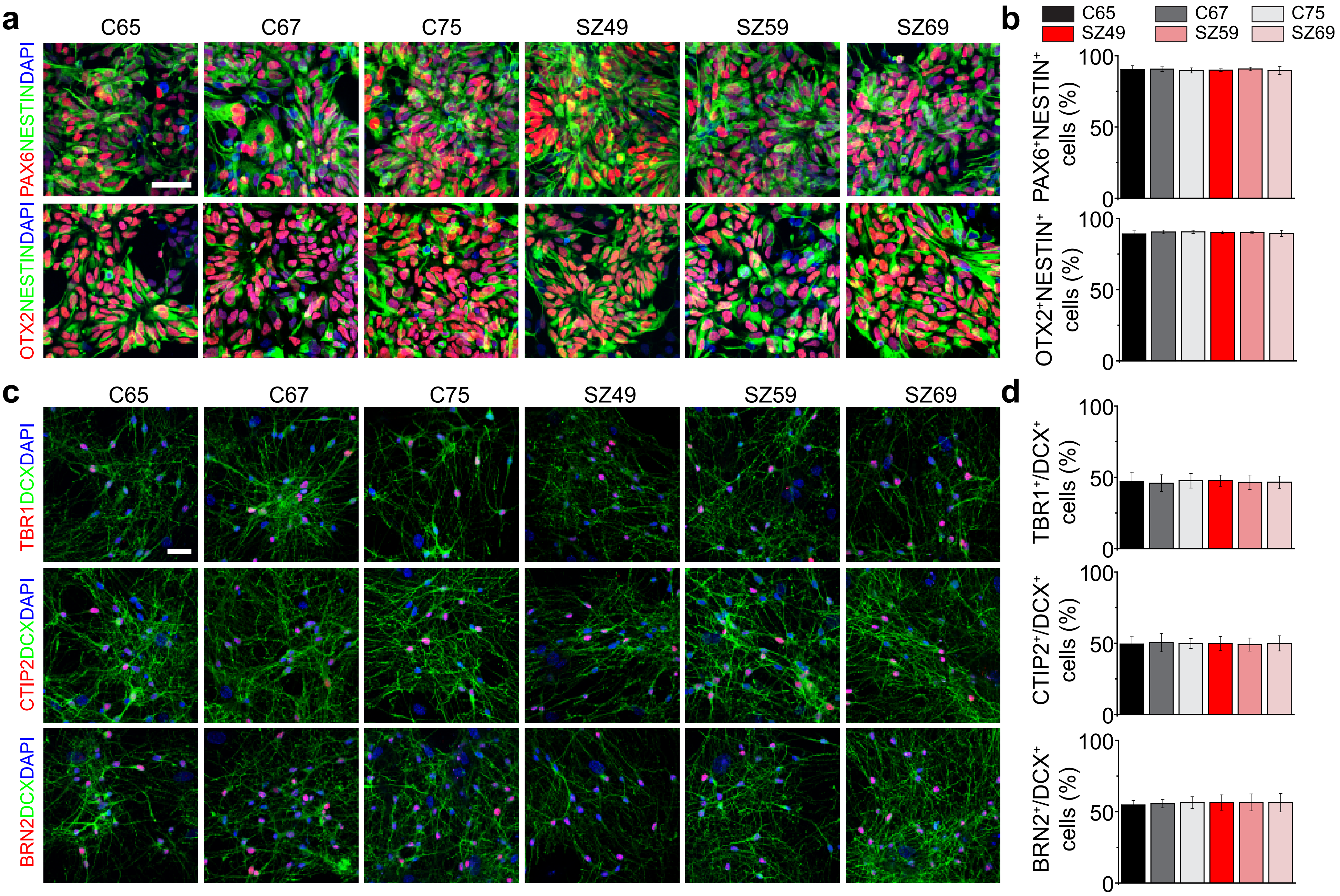


Supplementary Figure 9 (Guo et al.,)

### Supplemental Figure 10

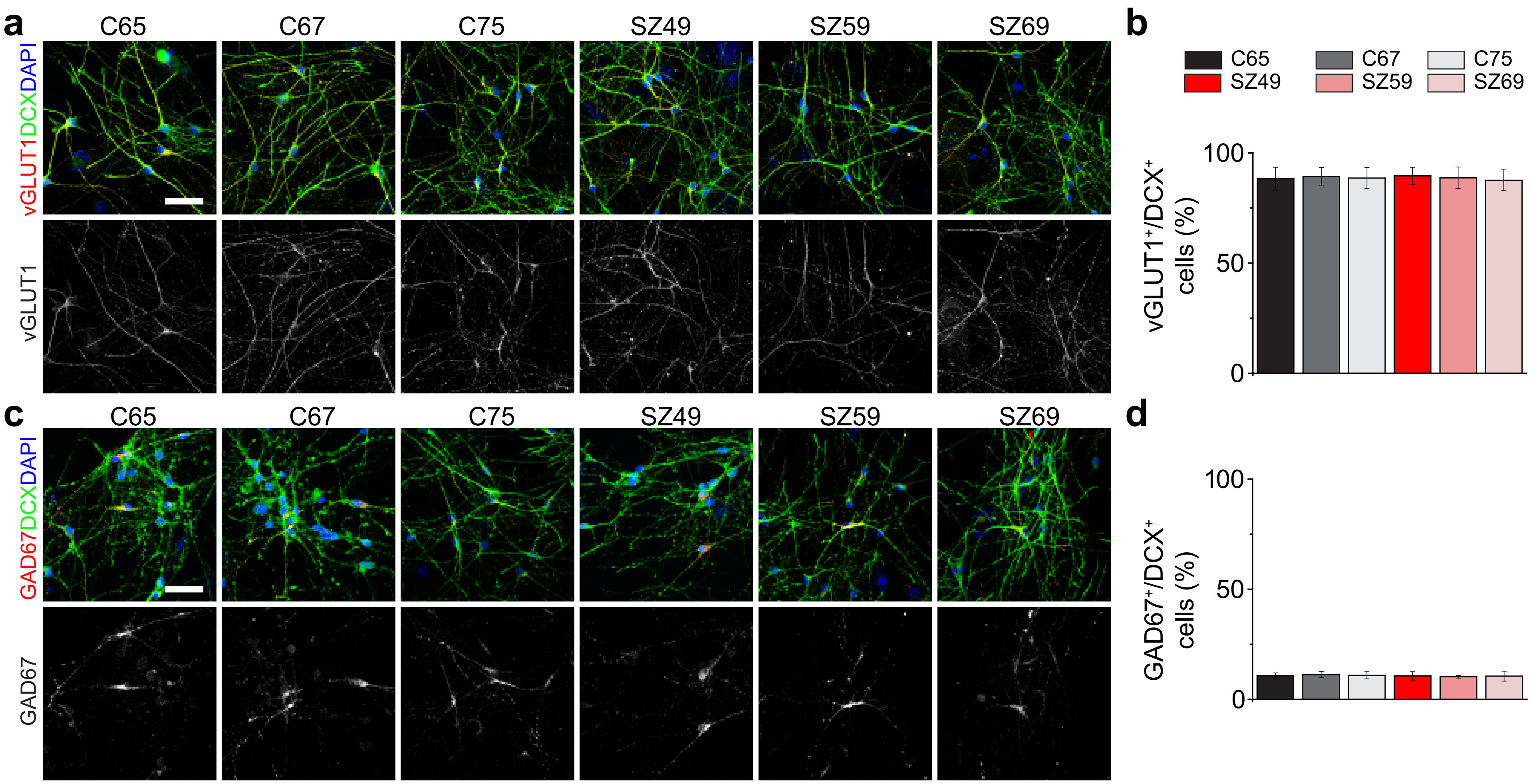


Supplementary Figure 10 (Guo et al.,)

### Supplemental Figure 11

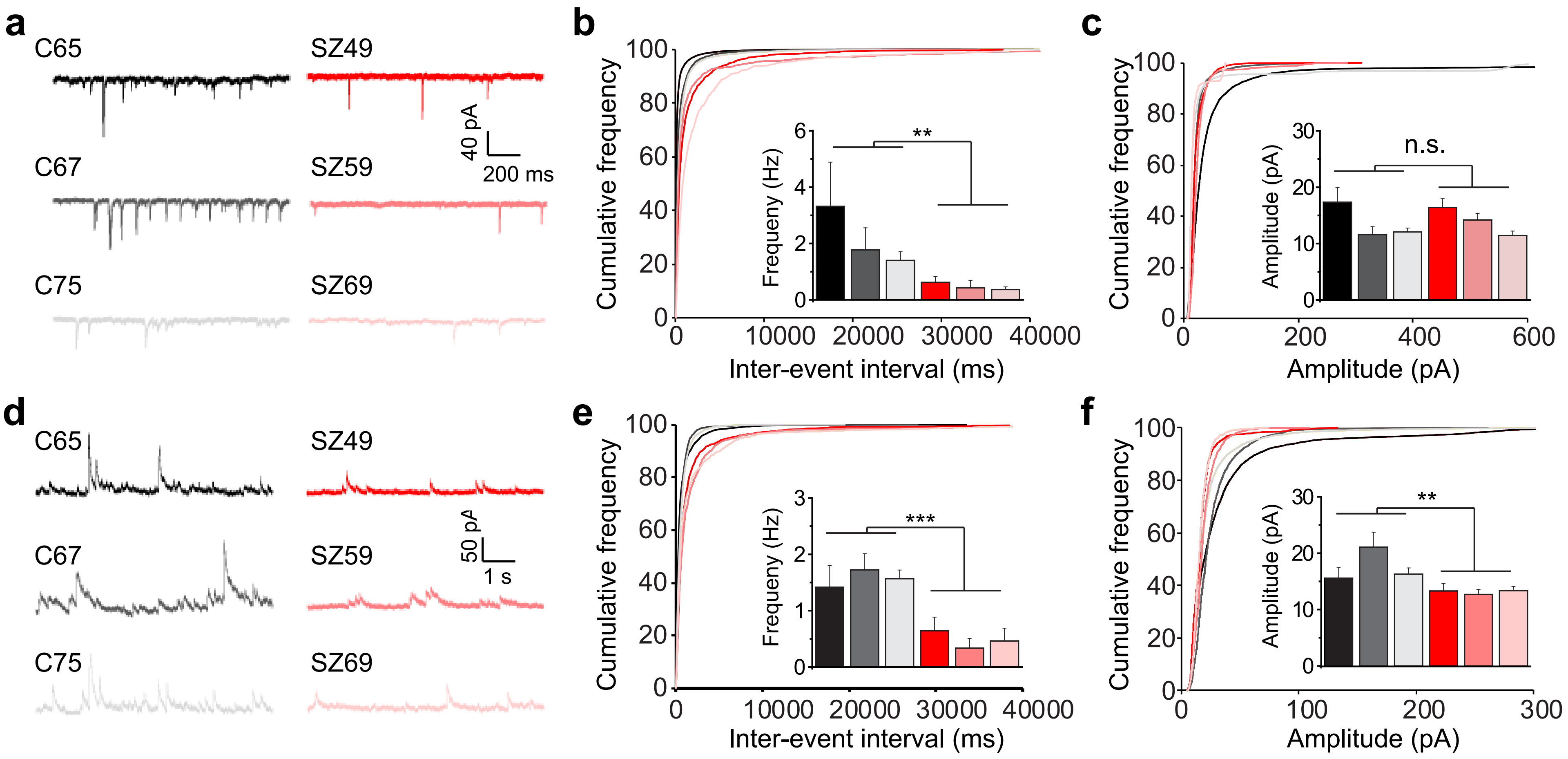


Supplementary Figure 11 (Guo et al.,)

### Supplemental Figure 12

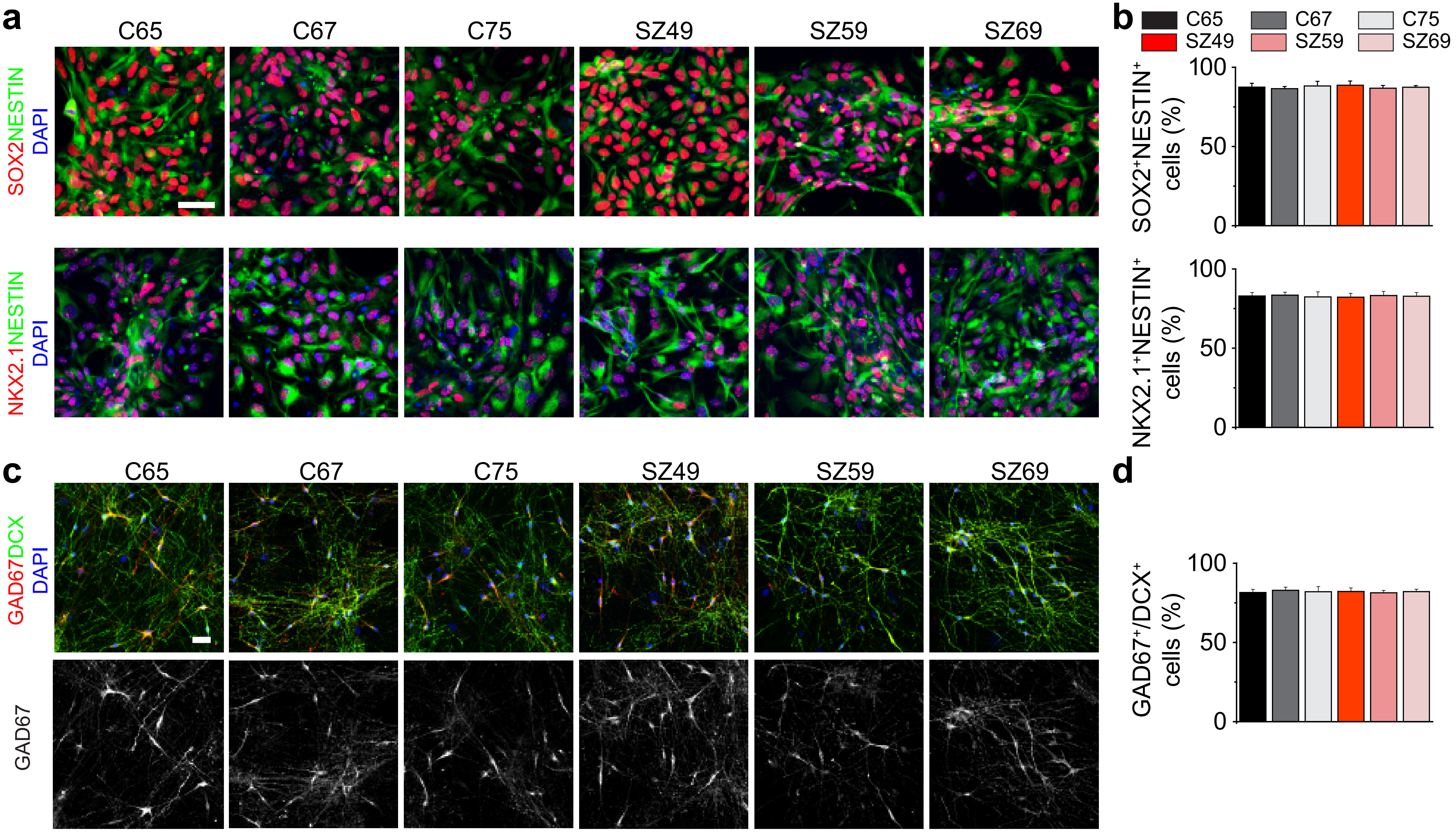


Supplementary Figure 12 (Guo et al.,)
