## Supplemental Figure 13 for "GABAergic neuron dysregulation in a human neurodevelopmental model for major psychiatric disorders"

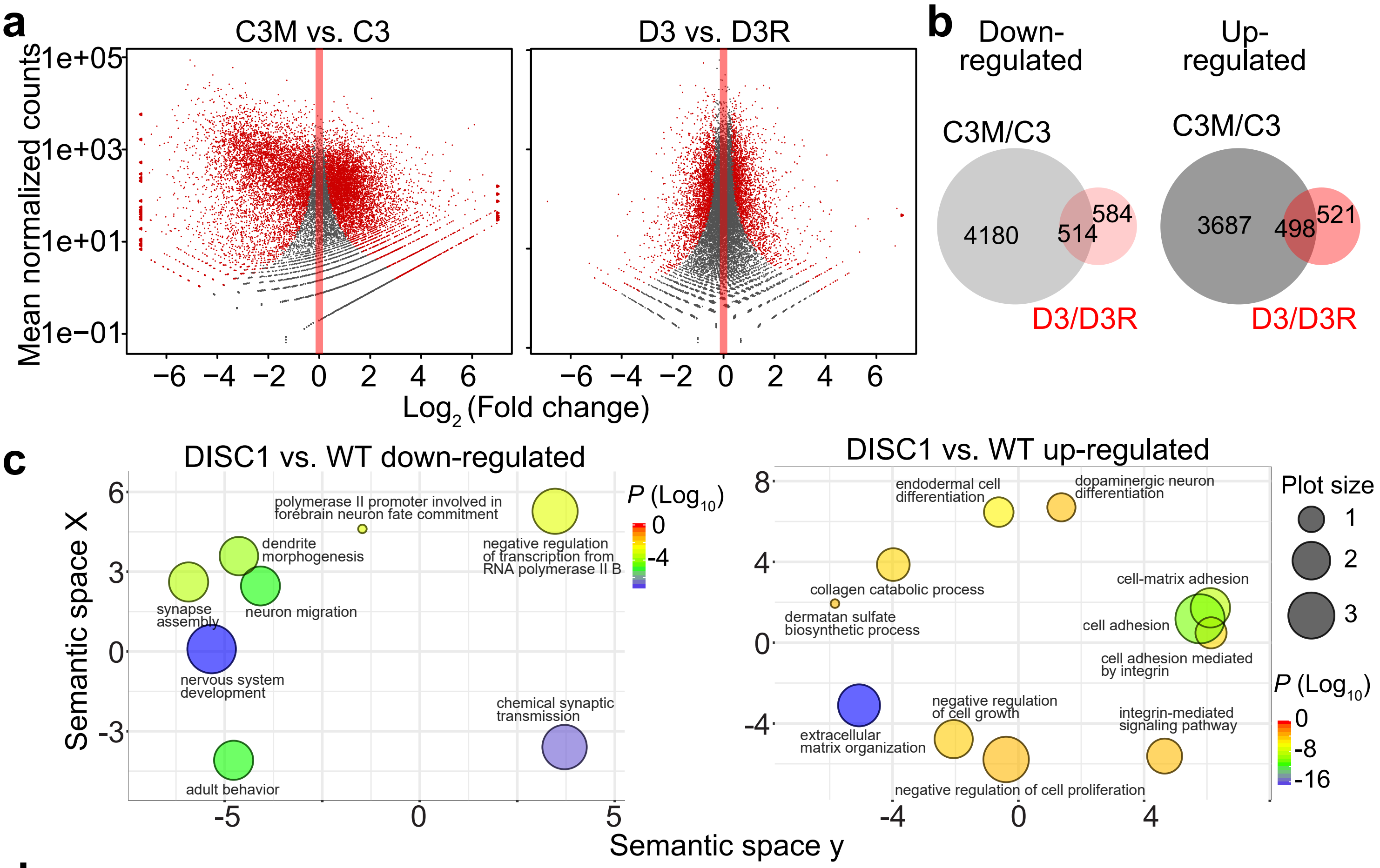


d

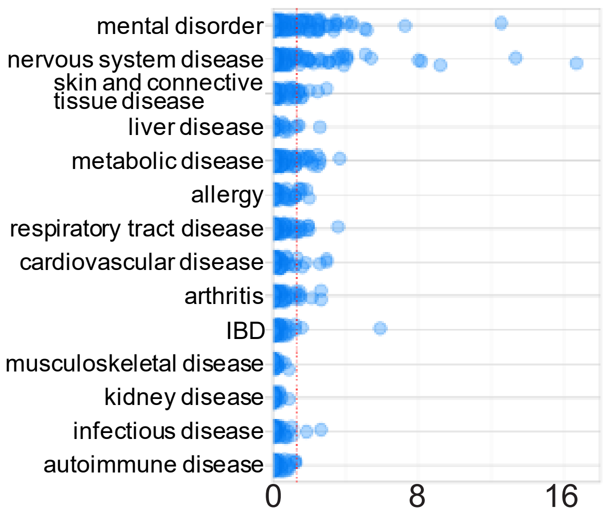


| Disease Category | # Studies | # $P < 0.05$ | BIONOM $P$ Value |
|---|---|---|---|
| mental disorder | 232 | 58 | 2.9E-09 |
| nervous system disease | 210 | 38 | 0.002 |
| skin and connective tissue disease | 78 | 10 | 0.279 |
| liver disease | 26 | 3 | 0.348 |
| metabolic disease | 154 | 16 | 0.603 |
| allergy;respiratory tract disease | 119 | 10 | 0.815 |
| cardiovascular disease | 92 | 6 | 0.914 |
| arthritis | 103 | 7 | 0.915 |
| inflammatory bowel disease (IBD) | 68 | 2 | 0.988 |
| musculoskeletal disease | 37 | 0 | 0.989 |
| kidney disease | 43 | 0 | 0.995 |
| infectious disease | 87 | 2 | 0.998 |
| autoimmune disease | 77 | 0 | 1.000 |
| **TOTAL** | **1326** | **152** | |

e

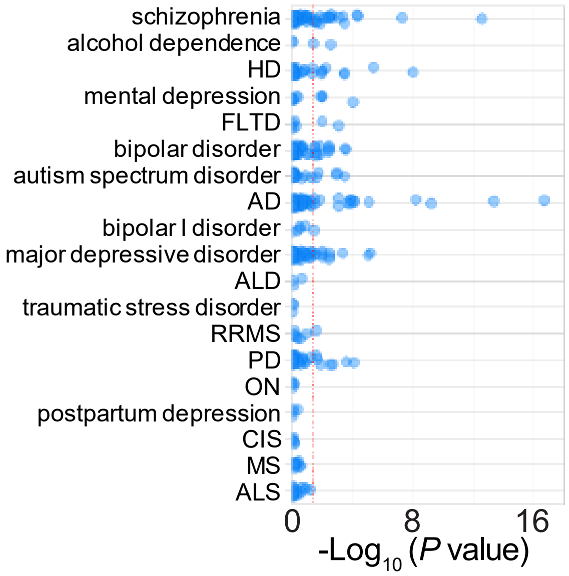


| Disease State | # Studies | # $P < 0.05$ | BIONOM $P$ Value |
|---|---|---|---|
| schizophrenia | 63 | 20 | 0.037 |
| alcohol dependence | 4 | 2 | 0.040 |
| Huntington's disease (HD) | 29 | 10 | 0.049 |
| mental depression | 11 | 4 | 0.083 |
| frontotemporal lobar degeneration (FTLD) | 7 | 2 | 0.200 |
| bipolar disorder | 49 | 13 | 0.213 |
| autism spectrum disorder | 21 | 5 | 0.343 |
| Alzheimer's disease (AD) | 68 | 16 | 0.381 |
| bipolar I disorder | 6 | 1 | 0.414 |
| major depressive disorder | 55 | 12 | 0.497 |
| adrenoleukodystrophy (ALD) | 3 | 0 | 0.541 |
| posttraumatic stress disorder | 3 | 0 | 0.541 |
| relapsing-remitting MS (RRMS) | 8 | 1 | 0.577 |
| Parkinson's disease (PD) | 36 | 7 | 0.600 |
| optic neuritis (ON) | 4 | 0 | 0.646 |
| postpartum depression | 4 | 0 | 0.646 |
| clinically isolated syndrome (CIS) | 5 | 0 | 0.727 |
| multiple sclerosis (MS) | 11 | 0 | 0.942 |
| amyotrophic lateral sclerosis (ALS) | 20 | 0 | 0.994 |
| **TOTAL** | **407** | **93** | |

Supplementary Figure 13 (Guo et al.,)
