## Supplemental Figure 14 for "GABAergic neuron dysregulation in a human neurodevelopmental model for major psychiatric disorders"

**a**

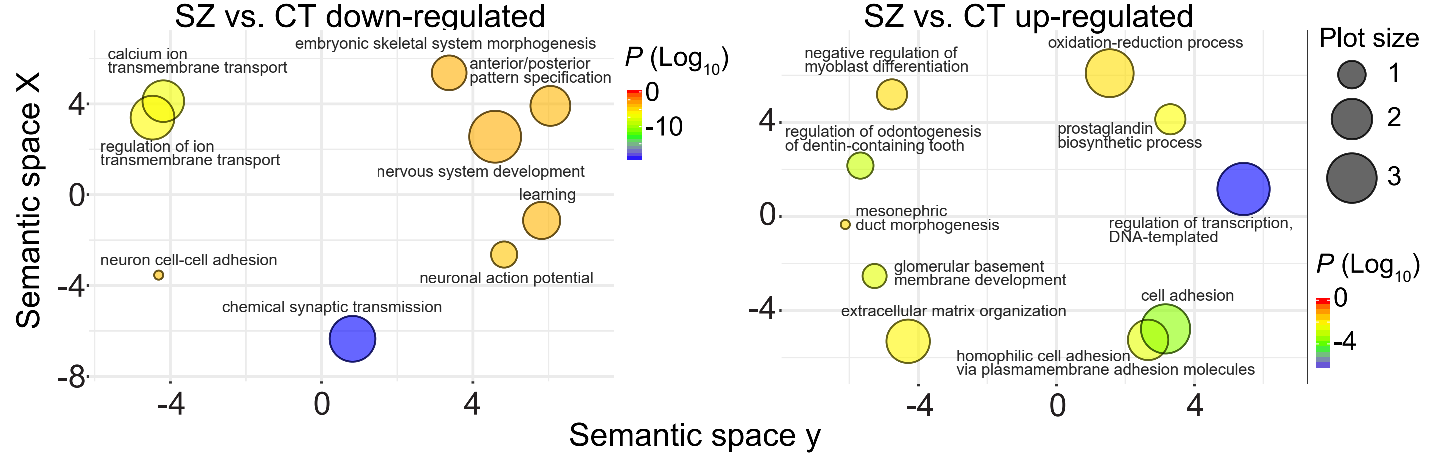


**b**

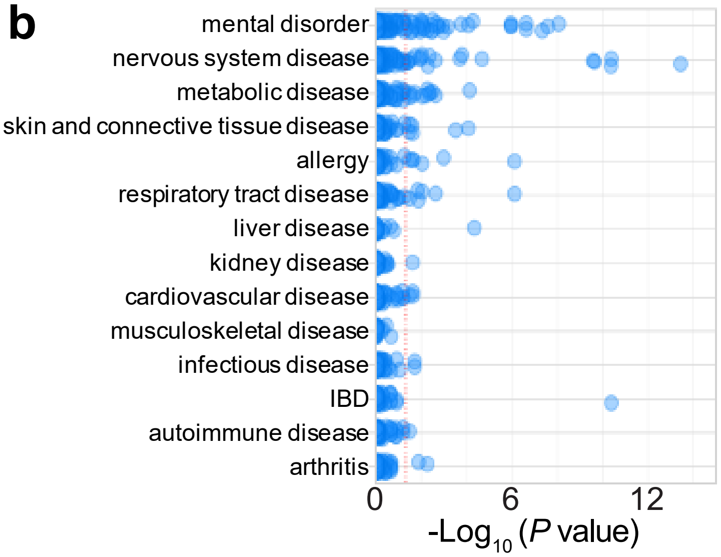


| Disease Category | # Studies | # $P < 0.05$ | BIONOM *P* Value |
|---|---|---|---|
| mental disorder | 232 | 37 | 1.3E-06 |
| nervous system disease | 210 | 19 | 0.110 |
| metabolic disease | 154 | 12 | 0.298 |
| skin and connective tissue disease | 78 | 6 | 0.316 |
| allergy;respiratory tract disease | 119 | 9 | 0.336 |
| liver disease | 26 | 1 | 0.559 |
| kidney disease | 43 | 1 | 0.819 |
| cardiovascular disease | 92 | 3 | 0.898 |
| musculoskeletal disease | 37 | 0 | 0.934 |
| infectious disease | 87 | 2 | 0.951 |
| inflammatory bowel disease (IBD) | 68 | 1 | 0.958 |
| autoimmune disease | 77 | 1 | 0.976 |
| arthritis | 103 | 2 | 0.980 |
| **TOTAL** | **1326** | **94** | |

**c**

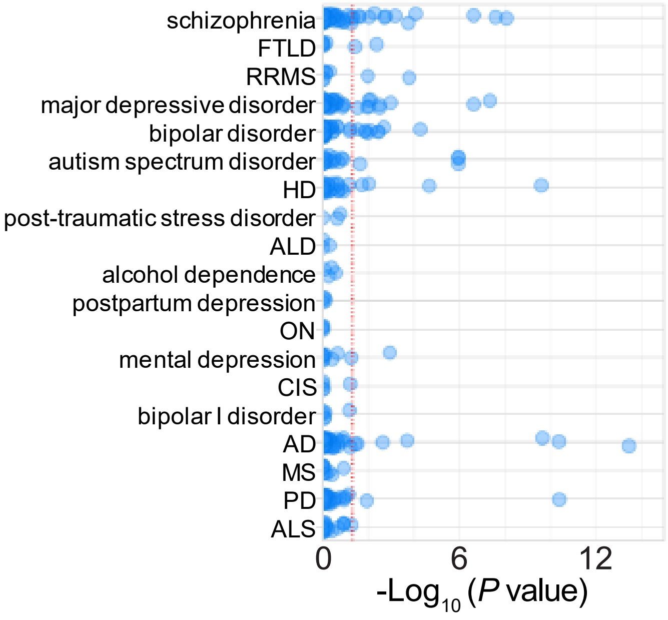


| Disease State | # Studies | # $P < 0.05$ | BIONOM *P* Value |
|---|---|---|---|
| schizophrenia | 63 | 13 | 0.035 |
| frontotemporal lobar degeneration (FTLD) | 7 | 2 | 0.054 |
| relapsing-remitting MS (RRMS) | 8 | 2 | 0.078 |
| major depressive disorder | 55 | 10 | 0.105 |
| bipolar disorder | 49 | 9 | 0.107 |
| autism spectrum disorder | 21 | 4 | 0.136 |
| Huntington's disease (HD) | 29 | 4 | 0.339 |
| post-traumatic stress disorder | 3 | 0 | 0.348 |
| adrenoleukodystrophy (ALD) | 3 | 0 | 0.348 |
| alcohol dependence | 4 | 0 | 0.434 |
| postpartum depression | 4 | 0 | 0.434 |
| optic neuritis (ON) | 4 | 0 | 0.434 |
| mental depression | 11 | 1 | 0.440 |
| clinically isolated syndrome (CIS) | 5 | 0 | 0.509 |
| bipolar I disorder | 6 | 0 | 0.574 |
| Alzheimer's disease (AD) | 68 | 7 | 0.696 |
| multiple sclerosis (MS) | 11 | 0 | 0.791 |
| Parkinson's disease (PD) | 36 | 2 | 0.874 |
| amyotrophic lateral sclerosis (ALS) | 20 | 0 | 0.942 |
| **TOTAL** | **407** | **54** | |

Supplementary Figure 14 (Guo et al.,)
